# Ancient Fennoscandian genomes reveal origin and spread of Siberian ancestry in Europe

**DOI:** 10.1101/285437

**Authors:** Thiseas C. Lamnidis, Kerttu Majander, Choongwon Jeong, Elina Salmela, Anna Wessman, Vyacheslav Moiseyev, Valery Khartanovich, Oleg Balanovsky, Matthias Ongyerth, Antje Weihmann, Antti Sajantila, Janet Kelso, Svante Pääbo, Päivi Onkamo, Wolfgang Haak, Johannes Krause, Stephan Schiffels

## Abstract

European history has been shaped by migrations of people, and their subsequent admixture. Recently, evidence from ancient DNA has brought new insights into migration events that could be linked to the advent of agriculture, and possibly to the spread of Indo-European languages. However, little is known so far about the ancient population history of north-eastern Europe, in particular about populations speaking Uralic languages, such as Finns and Saami. Here we analyse ancient genomic data from 11 individuals from Finland and Northwest Russia. We show that the specific genetic makeup of northern Europe traces back to migrations from Siberia that began at least 3,500 years ago. This ancestry was subsequently admixed into many modern populations in the region, in particular populations speaking Uralic languages today. In addition, we show that ancestors of modern Saami inhabited a larger territory during the Iron Age than today, which adds to historical and linguistic evidence for the population history of Finland.

## Introduction

The genetic structure of Europeans today is the result of several layers of migration and subsequent admixture. The incoming source populations no longer exist in unadmixed form, but have been identified using ancient DNA in several studies over the last few years^1–8^. Broadly, present-day Europeans have ancestors in three deeply diverged source populations: European hunter-gatherers who settled the continent in the Upper Paleolithic, Europe’s first farmers who expanded from Anatolia across Europe in the early Neolithic starting around 8,000 years ago, and groups from the Pontic Steppe that arrived in Europe during the final Neolithic and early Bronze Age ∼4,500 years ago. As a consequence, most Europeans can be modelled as a mixture of these three ancestral populations^3^.

The model of three ancestral populations, however, does not fit well for the present-day populations from north-eastern Europe such as Saami, Russians, Mordovians, Chuvash and Finns: they carry an additional ancestry component seen as increased allele sharing with modern East Asian populations^1,3,9,10^. The origin and timing of this East Asian-related contribution is unknown. Modern Finns are known to possess a distinct genetic structure among today’s European populations^9,11–14^, and the country’s geographical location at the crossroads of eastern and western influences introduces a unique opportunity to investigate the migratory past of North-East Europe. Furthermore, the early migrations and the genetic origins of the Saami people in relation to the Finnish population call for a closer inspection, with the linguistic evidence suggesting that Saami languages have dominated the whole of the Finnish region before 1,000 CE^15–17^. Here, the early Metal Age, Iron Age, and historical burials analysed provide a suitable time-transect to ascertain the timing of the arrival of the deeply rooted Siberian genetic ancestry, and a frame of reference for investigating linguistic diversity in the region today.

In this study we present new genome-wide data from 11 individuals from Finland and the Russian Kola Peninsula who lived between 3,500 to 200 years ago. In addition, we present a new high-coverage genome from a modern Saami individual for whom low coverage data was published previously^1^. Our results suggest that a new genetic component with strong Siberian affinity first arrived in Europe around 4,000 years ago, as observed in our oldest analysed individuals from northern Russia, and that the gene pool of modern north-eastern Europeans in general, and speakers of Uralic languages in particular, is the result of multiple admixture events between Eastern and Western sources since that first appearance. Additionally, we gain further insights into the genetic history of the Saami in Finland, by showing that during the Iron Age, close genetic relatives of modern Saami lived in an area much further south than their current geographic range.

## Results

### Sample information and archaeological background

The ancient individuals analysed in this study come from three time periods (Table 1, Figure 1, Supplementary Text 1). The six early Metal Age individuals were obtained from an archaeological site at Bolshoy Oleni Ostrov in the Murmansk Region on the Kola Peninsula (“Bolshoy” from here on). The site has been radiocarbon dated to 1610-1436 calBCE and the mitochondrial DNA HVR-I haplotypes from these six individuals have been previously reported^18^. Today, the region is inhabited by Saami. Seven individuals stem from excavations in Levänluhta, a lake burial in Isokyrö, Finland. Artefacts from the site have been dated to the Finnish Iron Age (400-800 CE). Today, the inhabitants of the area speak Finnish and Swedish. Two individuals were obtained from the 18-19^th^ century Saami cemetery of Chalmny Varre on the Russian Kola Peninsula. The cemetery and the surrounding area were abandoned in the 1960s because of planned industrial constructions, and hence became the subject of excavations. In addition, we sequenced the whole genome of a modern Saami individual to 17.5-fold coverage, for whom genotyping data has previously been published^1^.

**Table 1.**
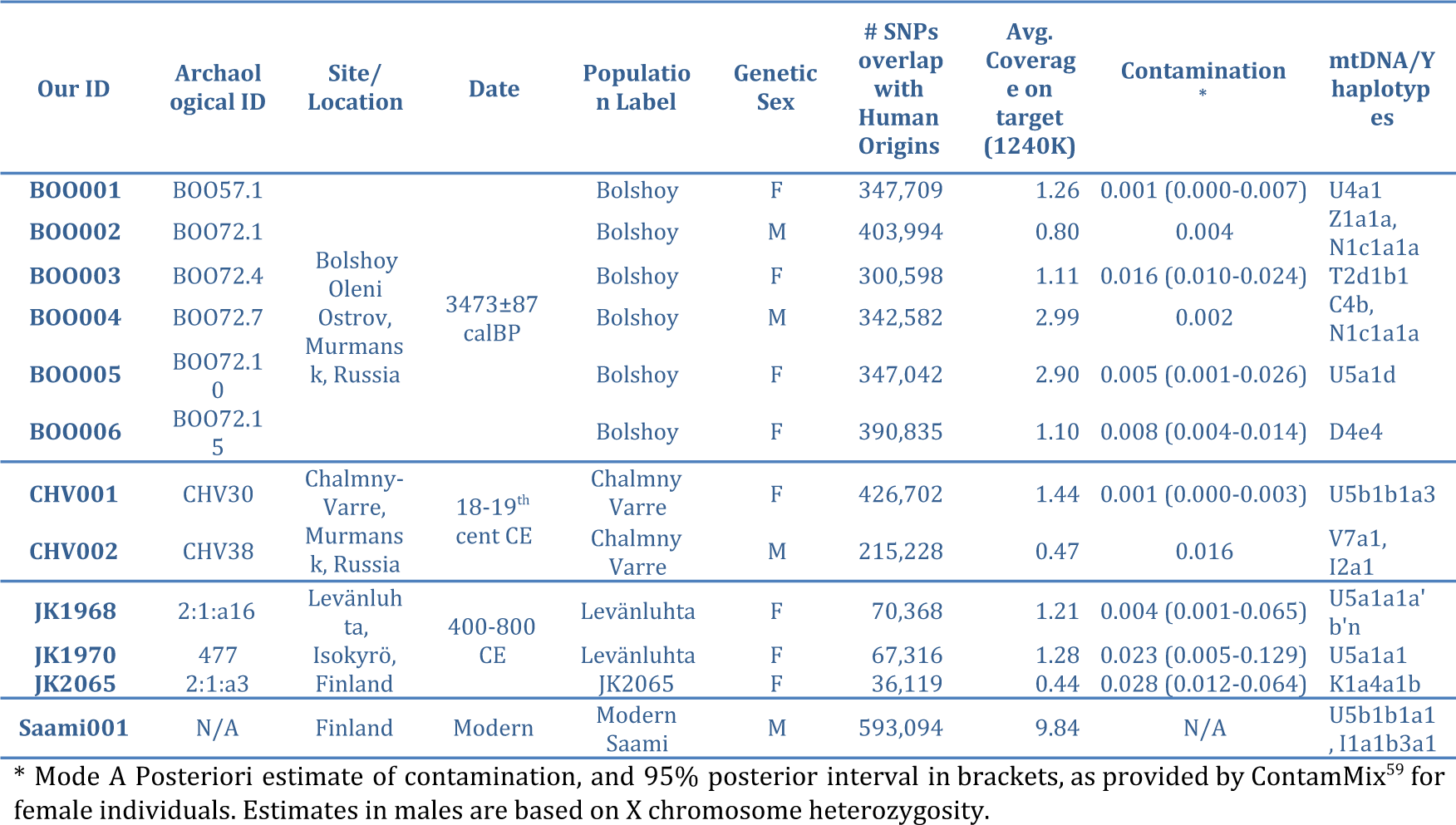
Sample Information

**Figure 1.**
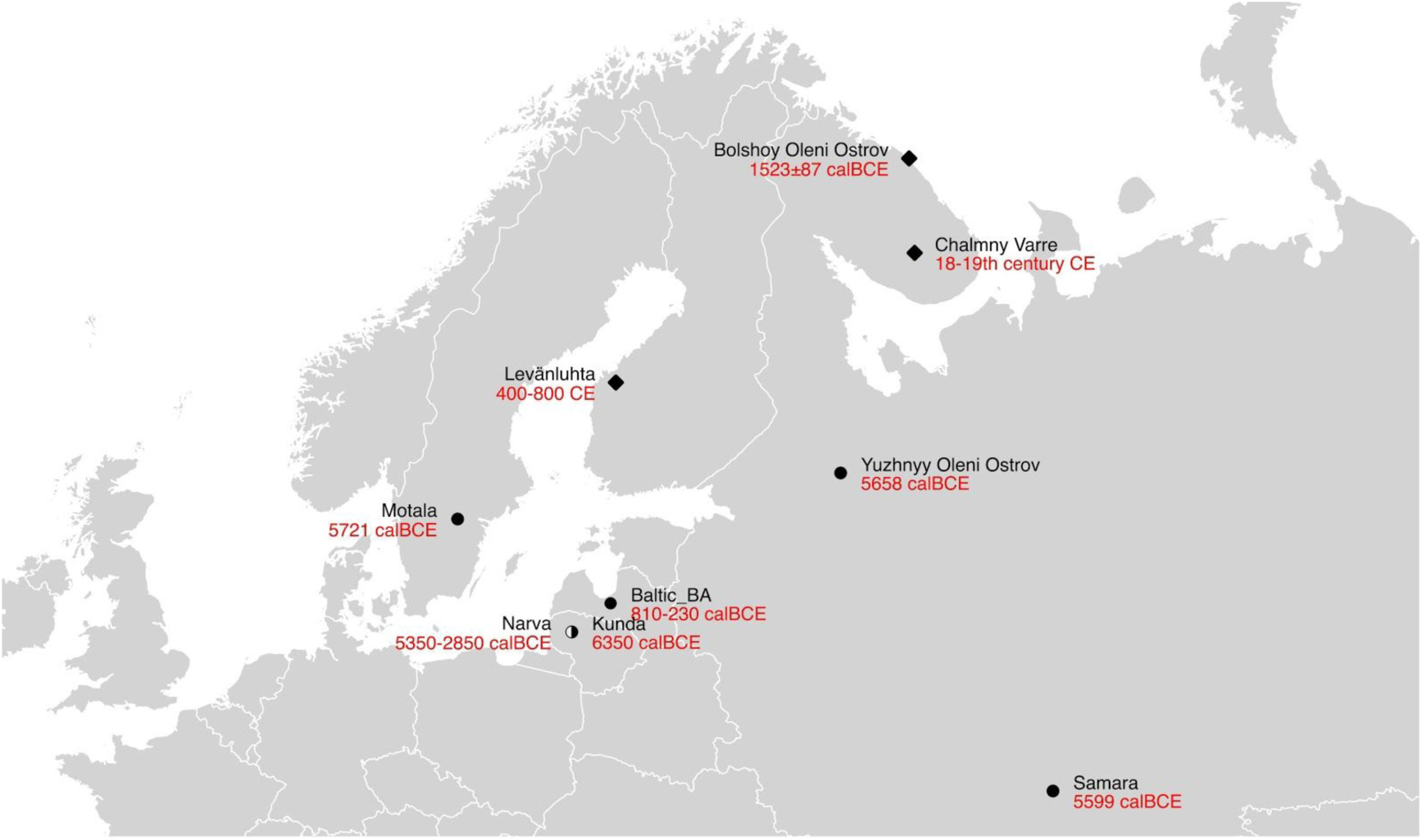
Location and age of archaeological sites with material used in this study (diamonds), and other sites relevant to this study (circles).

The sampling and subsequent processing of the ancient human remains was done in dedicated clean room facilities (Materials and Methods). A SNP-capture approach targeting a set of 1,237,207 single nucleotide polymorphisms (SNPs) was used to enrich our ancient-DNA libraries for human DNA^4^. The sequenced DNA fragments were mapped to the human reference genome, and pseudohaploid genotypes were called based on a random read covering each targeted SNP (see Materials and Methods). To ensure the ancient origin of our samples, and the reliability of the data produced, we implemented multiple quality controls. First, we looked at deamination patterns in the terminal bases of DNA reads, characteristic of ancient DNA (1.04%?4.50% in the 5’ end for UDG treated libraries) (Supplementary Table 1). Second, we estimated potential contamination rates through heterozygosity on the single-copy X chromosome for male individuals (all below 1.6%)^19^ (Table 1, see Supplementary Figure 1 for sex determination). Third, we computed statistics of the form *f*_*4*_(All reads, Damaged reads; Contamination source, Chimp), for six divergent modern populations to inspect the possible contamination sources (Supplementary Table 2). Fourth, we carried out supervised genetic clustering using ADMIXTURE^20^, using the same six divergent populations as defined clusters, to identify genetic disimilarities and possible contamination (Supplementary Figure 2). Finally, for individuals with sufficient SNP coverage, we carried out principal component analysis, projecting PMD-filtered and non-filtered versions of each individual to show that the different databases cluster together regardless of PMD-filtering^21^ (see Methods). Eleven ancient individuals passed the quality control thresholds, while four individuals were excluded from further analyses, due to low SNP coverage (< 15,000 SNPs). We merged these 11 individuals with 3,333 published present-day individuals genotyped on the Affymetrix Human Origins platform and 511 ancient individuals sequenced using a mixture of DNA capture and shotgun sequencing. ^1–4,8,22,23^.

**Figure 2.**
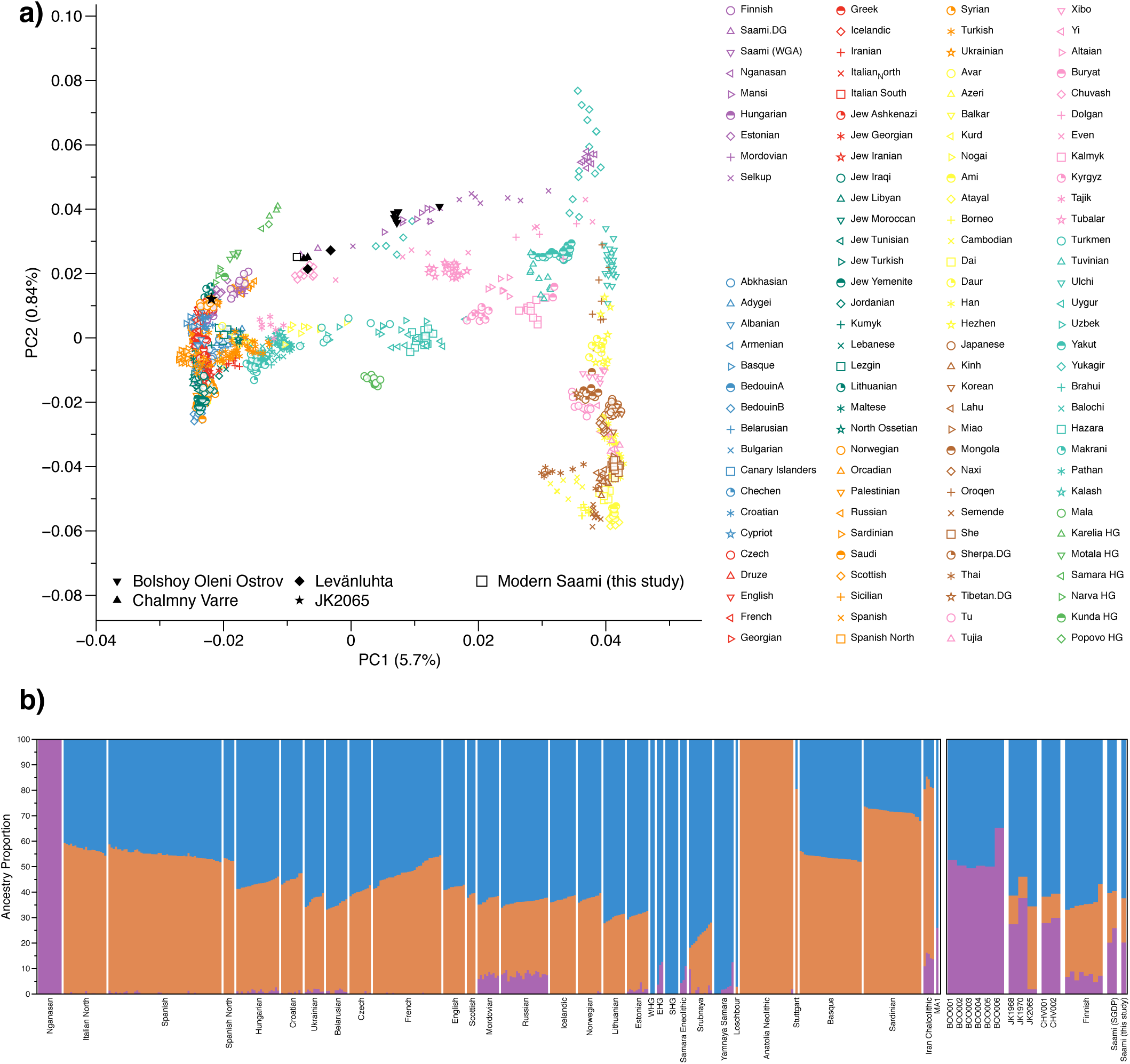
a) PCA plot of 113 Modern Eurasian populations, with individuals from this study projected on the principal components. Uralic speakers are highlighted in light purple. PCA of Europe can be found in Supplementary Figure 3. b) Plot of ADMIXTURE (K=3) results containing West Eurasian populations and the Nganasan. Ancient individuals from this study are represented by thicker bars.

### Eastern genetic affinities in Northern Europe

To investigate the genetic affinities of the individuals sampled, we projected them onto principal components computed from 1,320 modern European and Asian individuals (Figure 2a, Supplementary Figures 3a&b for a version focusing on West Eurasia). As expected, PC1 separates East Asian from West Eurasian populations. Within each continental group, genetic variability is spread across PC2: The East Asian genetic cline spans populations between the Siberian Nganasan (Uralic speakers) and Yukagirs at one end, and the Ami and Atayal from Taiwan at the other end. The West Eurasian cline along PC2 spans from the Bedouins on the Arabian Peninsula to north-eastern Europeans including Lithuanians, Norwegians and Finns. Between these two main Eurasian clines exist multiple clines, spanning between West and East Eurasians. These clines are likely the result of admixture events and population movements between East and West Eurasia. Most relevant to the populations analysed here is the admixture cline between north-eastern Europe and the North Siberian Nganasan, including mostly Uralic-speaking populations in our dataset (marked in light purple in Figure 2a).

**Figure 3.**
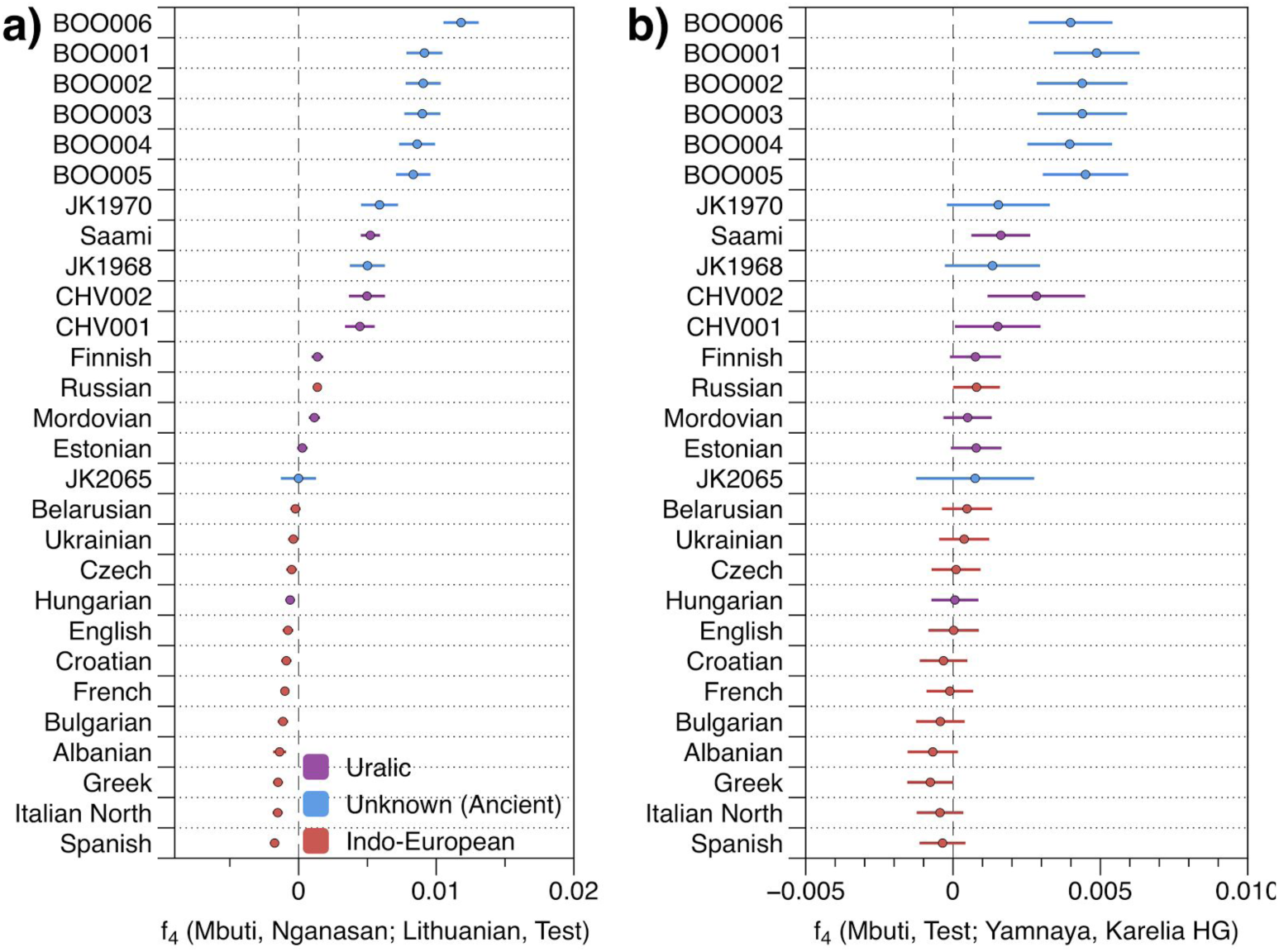
a) Calculated f_4_ (Mbuti, Nganasan; Lithuanian, Test). b) f_4_ (Mbuti, Test; Yamnaya, Karelia HG). Error bars represent 3 standard errors. Populations are coloured based on language family. Ancient individuals with unknown language are shown in blue.

Ten of the eleven ancient individuals from this study fall on this Uralic cline, with the exception of one individual from Levänluhta (JK2065), who is instead projected closer to modern Lithuanian, Norwegian and Icelandic populations. Specifically, two Levänluhta individuals and the two historical Saami from Russia are projected very close to two previously published modern Saami (Saami.DG)^23^ and the new Saami genome generated in this study (as well as the previously published genome of the same individual, here labelled “Saami (WGA)”^1^), suggesting genetic continuity in the North from the Iron Age to modern-day Saami populations. In contrast, the six ancient individuals from Bolshoy are projected much further towards East Asian populations, in an intermediate position along the Uralic cline and close to modern-day Mansi.

Unsupervised genetic clustering analysis as implemented in the ADMIXTURE^20^ program suggests a similar profile: north-eastern European populations harbour a Siberian genetic component (light purple) maximized in the Nganasan (Figure 2b). The hunter-gatherer genetic ancestry in Europeans (blue) is maximized in European Upper Palaeolithic hunter-gatherers, including the 8,000-year-old Western European hunter-gatherers from Hungary and Spain (WHG), the 8,000-year-old Scandinavian hunter-gatherers from Motala (SHG) and the 7,000-year-old individuals from Karelia in north-eastern Europe (EHG). An ancestry component associated with Europe’s first farmers (orange) is maximized in ancient individuals from Neolithic Anatolia. Within modern Europeans, the Siberian genetic component is maximized in the Saami, and can also be seen in similar proportions in the historical Saami from Chalmny Varre and in two of the Levänluhta individuals. The third Levänluhta individual, JK2065, falling also in an outlier position on the PCA, lacks the Siberian component. The six ancient individuals from Bolshoy show substantially higher proportions of the Siberian component, which comprises about half of their ancestry (49.4-65.3 %), whereas the older Mesolithic individuals from Motala do not share this Siberian ancestry. The Siberian ancestry seen in EHG probably corresponds to a previously reported affinity towards Ancient North Eurasians (ANE)^2,24^, which also comprises part of the ancestry of Nganasans. Interestingly, results from uniparentally-inherited markers (mtDNA and Y chromosome) as well as certain phenotypic SNPs also show Siberian signals in Bolshoy: mtDNA haplogroups Z1, C4 and D4, common in modern Siberia^18,25,26^, in individuals BOO002, BOO004 and BOO006, respectively (confirming previous findings^18^), as well as Y-chromosomal haplotype N1c1a1a (N-L392) in individuals BOO002 and BOO004. Haplogroup N1c, to which this haplotype belongs, is the major Y chromosomal lineage in modern North-East Europe and European Russia, especially in Uralic speakers, for example comprising as much as 54% of Eastern Finnish male lineages today^27^. Notably, this is the earliest known occurrence of Y-haplogroup N1c in Fennoscandia. Additionally, within the Bolshoy population we observe the derived allele of rs3827760 in the *EDAR* gene, which is found in near-fixation in East Asian and Native American populations but is extremely rare elsewhere^28^, and has been linked to phenotypes related to tooth shape^29^ and hair morphology^30,31^ (Supplementary Table 7). Scandinavian hunter-gatherers from Motala in Sweden have also been found to carry haplotypes associated with this allele^4^. Finally, we see high frequencies of haplotypes associated with diets rich in high poly-unsaturated fatty acids, on the *FADS* genes^4,32,33^. The *FADS* haplotype observed here has previously been linked with Greenlanders^32^, and is found in lower frequencies within Europe^33^.

### The arrival of Siberian ancestry in Europe

We formally tested for admixture in north-eastern Europe by calculating *f*_*3*_(*Test*; Siberian source, European source) using Uralic-speaking populations -Estonians, Saami, Finnish, Mordovians and Hungarians -and Russians as *Test* populations. Significantly negative *f*_*3*_ values correspond to the *Test* population being admixed between populations related to the two source populations^34^. Additionally, the magnitude of the statistic is directly related to the ancestry composition of the tested source populations and how closely those ancestries are related to the actual source populations. We used multiple European and Siberian sources, to capture differences in ancestral composition among proxy populations. As proxies for the Siberian source we used Bolshoy, Mansi and Nganasan, and for the European source modern Icelandic, Norwegian, Lithuanian and French. Our results show that all of the test populations are indeed admixed, with the most negative values arising when Nganasan are used as the Siberian source (Supplementary Table 3). Among the European sources, Lithuanians gave the most negative results for Estonians, Russians and Mordovians. For modern Hungarians, the European source giving most negative results was French, while both Bolshoy and Nganasan gave equally negative results when used as the Siberian source. With Finns as test, modern Icelanders were the European source giving most negative statistics. Finally, the present-day Saami as a *Test* produced the most negative, though non-significant, result when Icelanders and Nganasan were used as the European and Siberian sources respectively. This non-significance is likely a result of the low number of modern Saami individuals in our dataset (n=3). To further test differential relatedness with Nganasan in European populations as well as the ancient individuals in this study, we calculated *f*_*4*_(Mbuti, Nganasan; Lithuanian, Test) (Figure 3a). Consistent with *f*_*3*_-statistics above, all the ancient individuals and modern Finns, Saami, Mordovians and Russians show excess allele sharing with Nganasan when used as *Test* populations. Of all Uralic speakers in Europe, Hungarians are the only population that shows no evidence of excess allele sharing with Nganasan, consistent with their distinct population history as evidenced by historical sources (see ref ^35^ and references therein).

We further estimated the amount of Siberian ancestry in these populations using *qpAdm*^*3*^. All ancient and modern individuals from the Baltics, Finland and Russia were successfully modelled as a mixture of five lines of ancestry, represented by Karelian EHG, Yamnaya from Samara, LBK from the early European Neolithic, western Mesolithic hunter-gatherers (from Spain, Luxembourg and Hungary), and Nganasan, or subsets of those five (Supplementary Table 4). Our motivation in including Karelian EHG as well as Yamnaya as sources in our model, although Yamnaya has been shown to contain large amounts of EHG ancestry^3^, is that the six Bolshoy individuals share significantly more alleles with Karelian EHG than with Yamnaya (Figure 3b). The 3,500-year-old ancient individuals from Bolshoy evidence the earliest presence, as well as the highest proportion of Siberian ancestry in this region (Figure 4). The geographically proximate ancient hunter-gatherers from the Baltics (8,000 and 8,300 BP) and Motala (∼8,000 BP), who predate Bolshoy, lack this component. In our models, Karelian EHG lack this component by construction, since they are one of the source populations. All later ancient individuals in this study have lower amounts of Siberian ancestry than Bolshoy, probably as a result of admixture with other populations from further south and dilution, as suggested by the increased proportion of early European farmer ancestry related to Neolithic Europeans (Figure 2b, Figure 3a).

**Figure 4.**
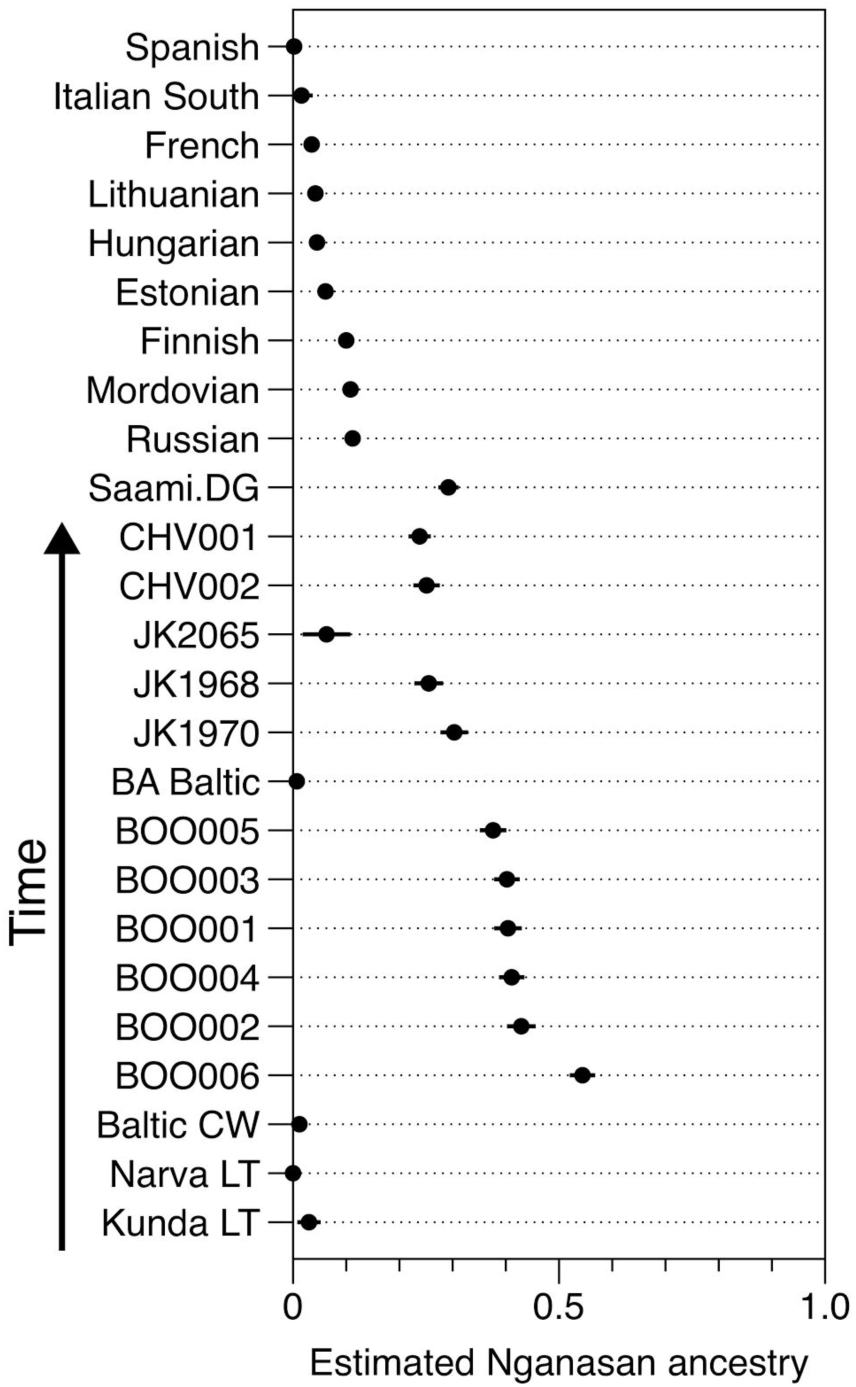
Estimated proportion of Nganasan-like ancestry, using qpAdm. Each Test population was explained as a 5-way mixture between Karelian HG, WHG, Yamnaya, LBK from the early Neolithic, and Nganasan. A set of seven outgroup populations (see Methods / Supplementary Table 4), which are differentially related to each of the five source populations, was used for all models. Results from only the least complex accepted model were used. The error bars represent one standard error around each reported proportion. The results for the Levänluhta samples rely only on transversions. CW=Corded Ware. LT=“from Lithuania”.

As shown by these multiple lines of evidence, the pattern of genetic ancestry observed in north-eastern Europe is the result of admixture between populations from Siberia and populations from Europe. To obtain a relative date of this admixture, and as an independent line of evidence thereof, we used admixture linkage disequilibrium decay, as implemented in ALDER^36^. ALDER provides a relative date estimate for a single-pulse admixture event in generations. When multiple admixture events have occurred, ALDER will provide a date that is an average across the admixture processes^36,37^. Based on the ALDER admixture date inferred for Bolshoy using Nganasan and EHG as admixture sources, and assuming a generation time of 29 years^38^, we estimate the time of introduction of Siberian ancestry in Bolshoy to 3,977 years before present (yBP) (Figure 5b). Estimates obtained using Nganasan and Lithuanians as source populations provided a similar estimate (Supplementary Figure 4 for LD decay plots for multiple populations using Lithuanians and Nganasan as sources.). However, for all other populations with evidence for Siberian ancestry, we find much younger admixture dates (Figure 5a), suggesting that the observed genetic ancestry in north-eastern Europe is inconsistent with a single-pulse admixture event. From the data shown in this study, the Bolshoy individuals mark the earliest evidence of Siberian ancestry in north-eastern Europe, while also predating the introduction of ancestry related to Neolithic farmers into this area. As such, the admixture date estimate provided by ALDER for Bolshoy is likely closer to the true time when Siberian ancestry was first introduced in the area, and not affected by later admixture events responsible for the introduction of central European ancestry into the region.

**Figure 5.**
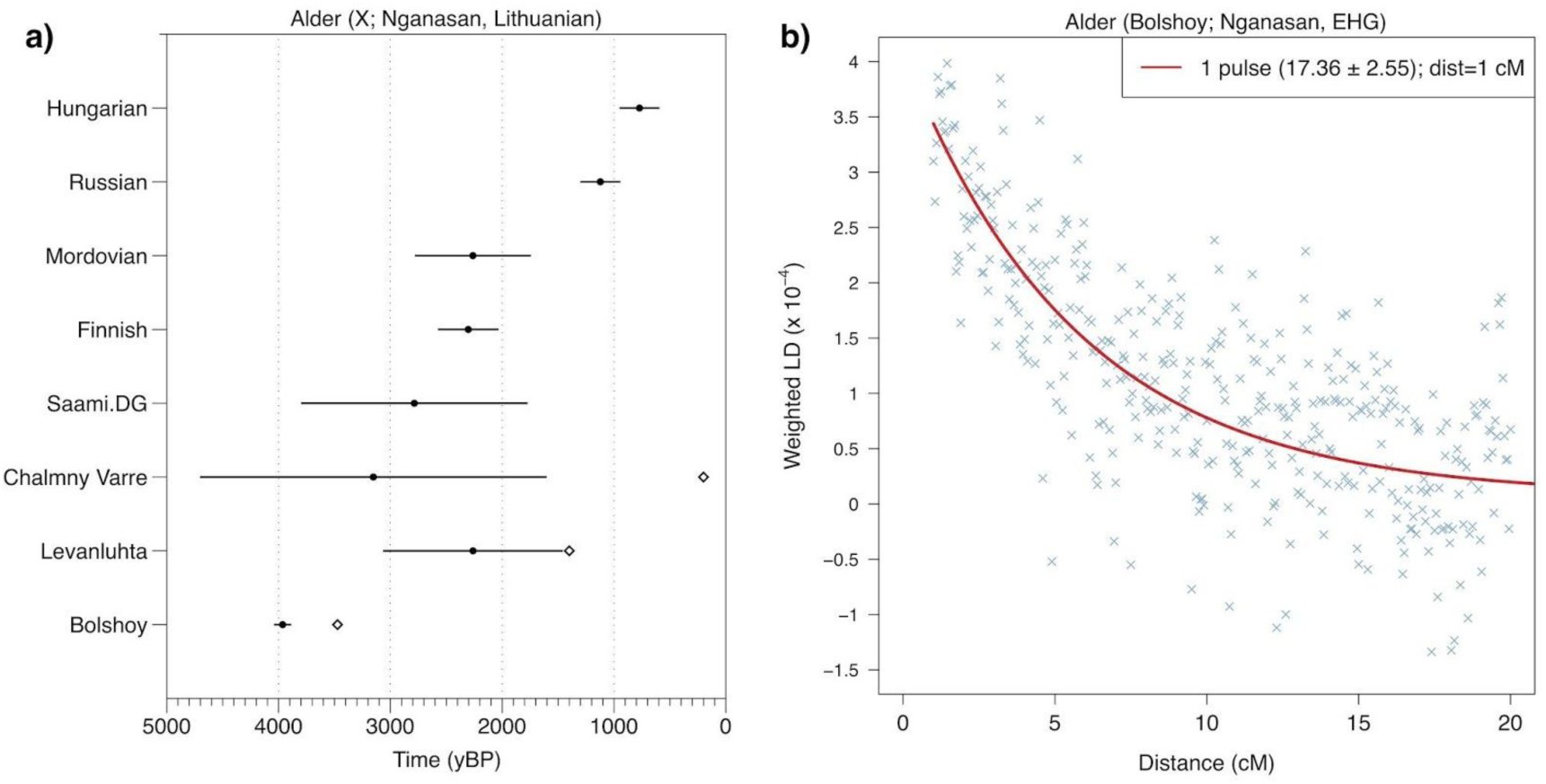
a) ALDER-inferred admixture dates (filled circles) for different populations, using Nganasan and Lithuanian as sources. Error bars represent errors provided by ALDER and include the uncertainty surrounding the dating of ancient population samples, calculated using standard propagation. Available dates for ancient populations are shown in white diamonds. b) LD decay curve for Bolshoy, using Nganasan and EHG as sources. The fitted trendline considers a minimum distance of 1cM. A full set of LD decay plots can be found in Supplementary Figure 4.

### Major genetic shift in Finland since the Iron Age

The population history of Finland is the subject of an ongoing discussion, with regard to the status of the Saami as the earlier inhabitants of Finland, compared to Finns. Many linguists claim that Saami languages were spoken in Finland prior to the arrival of the ancestors of modern Finns^15–17^. Particularly, southern Ostrobothnia, where Levänluhta is located, has been suggested to harbour a southern Saami dialect until the late first millennium, when early Finnish took over as the dominant language^39^. Historical sources note “Lapps” living in the parishes of central Finland still in the 1,500s^40^, though it is not entirely clear if all of them spoke Saami but some having been Finns who changed their means of livelihood from agriculture to hunting and fishing. There are also documents of intermarriage, although many of the indigenous people retreated to the north (see ref ^41^ and references therein). The ancestors of present-day Finns thus displaced but also admixed with the indigenous people of Finland, who are likely the ancestors of today’s Saami speakers^42–44^.

To test whether the ancestry of ancient individuals from Levänluhta is more closely related to modern-day Saami or to Finnish populations, we calculated *f*_4_(Saami(SGDP), *Test*; *X*, Mbuti) and *f*_4_(Finnish, *Test*; *X*, Mbuti), where *Test* was substituted with each ancient individual from Levänluhta, the two historical Saami individuals from Chalmny Varre, as well as the Modern Saami individual, and *X* was substituted by worldwide modern-day populations (Supplementary Tables 5 & 6, and Supplementary Figures 5 & 6). One Levänluhta and the two Chalmny Varre individuals consistently formed a clade with modern-day Saami, but not with modern-day Finns, with respect to all worldwide populations. One Levänluhta individual (JK1970) showed slightly lower affinity to central Europe than the modern-day Saami, while still rejecting a cladal position with modern-day Finns. This indicates that the people inhabiting Levänluhta during the Iron Age, and possibly other areas in the region as well, were more closely related to modern-day Saami than to present-day Finns.

One of the individuals from Levänluhta (JK2065) rejects a cladal position with modern Saami to the exclusion of most modern Eurasian populations. This individual also rejected a cladal position with Finns. By analysing four additional individuals with low coverage from the same site using PCA (Supplementary Figure 3), we confirm the outlier position of JK2065 compared to all other 6 individuals from that site, consistent also with the ADMIXTURE, and qpAdm results. The outlier position cannot be explained by modern contamination, since this individual passed several tests for authentication (see Methods) along with all other individuals. This individual therefore appears to be non-representative for the Levänluhta population, and evades confirmative interpretations.

## Discussion

In terms of ancient human DNA, north-eastern Europe has been relatively understudied. In this study we extend the available information from this area considerably, and present the first ancient genome-wide data from Finland. While the Siberian genetic component described here was previously described in modern-day populations from the region^1,3,9,10^, we gain further insights into its temporal depth. Our data suggest that this fourth genetic component found in modern-day north-eastern Europeans arrived in the area around 4,000 years ago at the latest, as illustrated by ALDER dating using the ancient genome-wide data from Bolshoy Oleni Ostrov. The upper bound for the introduction of this component is harder to estimate. The component is absent in the Karelian hunter-gatherers (EHG)^3^ dated to 8,300-7,200 yBP as well as Mesolithic and Neolithic populations from the Baltics from 8,300 yBP and 7,100-5,000 yBP respectively ^8^. While this suggests an upper bound of 5,000 yBP for the arrival of Siberian ancestry, we cannot exclude the possibility of its presence even earlier, yet restricted to more northern regions, as suggested by its absence in populations in the Baltic during the Bronze Age. Our study also presents the earliest occurrence of the Y-chromosomal haplogroup N1c in Fennoscandia. N1c is common among modern Uralic speakers, and has also been detected in Hungarian individuals dating to the 10^th^ century^35^, yet it is absent in all published Mesolithic genomes from Karelia and the Baltics^3,8,45,46^.

The large Siberian component in the Bolshoy individuals from the Kola Peninsula provides the earliest direct genetic evidence for an eastern migration into this region. Such contact is well documented in archaeology, with the introduction of asbestos-mixed Lovozero ceramics during the second millenium BC ^47^, and the spread of even-based arrowheads in Lapland from 1,900 BCE^48,49^. Additionally, the nearest counterparts of Vardøy ceramics, appearing in the area around 1,600-1,300 BCE, can be found on the Taymyr peninsula, much further to the east^48,49^. Finally, the Imiyakhtakhskaya culture from Yakutia spread to the Kola Peninsula during the same period^18,50^. Contacts between Siberia and Europe are also recognised in linguistics. The fact that the Siberian genetic component is consistently shared among Uralic-speaking populations, with the exceptions of Hungarians and the non-Uralic speaking Russians, would make it tempting to equate this component with the spread of Uralic languages in the area. However, such a model may be overly simplistic. First, the presence of the Siberian component on the Kola Peninsula at ca. 4000 yBP predates most linguistic estimates of the spread of Uralic languages to the area^51^. Second, as shown in our analyses, the admixture patterns found in historic and modern Uralic speakers are complex and in fact inconsistent with a single admixture event. Therefore, even if the Siberian genetic component partly spread alongside Uralic languages, it likely presented only an addition to populations carrying this component from earlier.

The novel genome-wide data here presented from ancient individuals from Finland opens new insights into Finnish population history. Two of the three higher coverage individuals and all six low coverage individuals from Levänluhta showed low genetic affinity to modern-day Finnish speakers of the area. Instead, an increased affinity was observed to modern-day Saami speakers, now mostly residing in the north of the Scandinavian Peninsula. These results suggest that the geographic range of the Saami extended further south in the past, and hints at a genetic shift at least in the western Finnish region during the Iron Age. The findings are in concordance with the noted linguistic shift from Saami languages to early Finnish. Further ancient DNA from Finland is needed to conclude to what extent these signals of migration and admixture are representative of Finland as a whole.

## Materials and Methods

### Sampling

Sampling and extracting ancient DNA requires a strict method for avoiding contamination introduced by contemporary genetic material. For 13 original Iron Age individuals from Finland, the sampling took place in a clean room facility dedicated to ancient DNA work, at the Institute for Archaeological Sciences in Tübingen. The procedure included documenting, photographing and storing the samples in individual, ID-coded plastic tubes and plastic bags. Since the specimens were teeth broken down in several fragments, the dentine was carefully detached from the enamel with a dentist drill and cooled-down diamond drill heads, rotated at the speed below 15 rpm, to avoid possible heat damage to the DNA.

For each specimen, ∼50mg of dentine powder was used for DNA extraction procedure specifically designed for ancient DNA material^52^. Altogether 100µl of extract was eluted in TET (10mM Tris-HCL, 1mM EDTA pH 0,8%, 0.1% Tween20). For samples from the sites of Bolshoy and Chalmny Varre we used leftover tooth powder that was originally processed at the Institute of Anthropology at the University of Mainz for replication purposes as described in Der Sarkissian et al. 2013.

### Radiocarbon date calibration

We calibrated the radiocarbon date of Bolshoy using Intcal13 as the calibration curve, using OxCal 4.3.

### DNA extraction and library preparation

DNA from Bolshoy and Chalmny Varre samples was extracted in the ancient DNA facilities of the Max-Planck Institute for the Science of Human History (MPI-SHH) in Jena, Germany. We used 50mg tooth powder for each DNA extraction using a modified version of the Dabney et al. 2013 protocol, resulting in 100µl of DNA extract. Negative controls were processed in parallel at a ratio of 1:7. Of the 100µl extract, 20µl of eluted DNA was immortalized as double-stranded library, following the Meyer and Kircher protocol^53^. Libraries were produced for all of the 13 extracts without a Uracil-DNA-glycosylase (UDG) treatment. The UDG treatment is normally introduced to eliminate the characteristic ancient DNA damage, which in non-treated library data is used to authenticate the aging process undergone by the sample. This makes the fragments and thus also the data retrieved more reliable for the use of population genetics. UDG-half treatment presents an intermediate option, where the damage is reduced to a minimum in the interior part of the DNA fragments, while still preserving a small indicative amount of deamination damage at the ends^54^.

Three of the original 13 libraries, estimated to pass a threshold of 1% endogenous human DNA after capture, were enriched with the 1240K SNP capture^4^, and later sequenced deeper for higher resolution. Additionally, for seven of the original 13 extracts from Levänluhta, and all Bolshoy and Chalmny Varre extracts, UDG-half treated libraries were produced, as described in^54^. For each library, a unique pair of 8 base pair long indexes was incorporated^55^. A positive control of cave bear DNA was used as bone powder in extractions, and as an extract for library preparation. Negative controls for both extraction and library preparation stages were kept alongside the samples throughout the entire workflow. All pre-amplification steps of the library preparation were conducted in the clean-room facilities of the University of Tübingen or the MPI-SHH in the case of BOO and CHV to minimize the risk of modern human contamination.

Experiment efficiency was ensured by quantifying the concentration of the libraries on qPCR (Roche) using aliquots from libraries before and after indexing. The positive controls confirmed the success of extraction and library preparation, whereas the negative controls showed 4-5 orders of magnitude lower concentration than samples, showing limited contamination levels from the laboratory processes.

The libraries were amplified with PCR, for the amount of cycles respective to the concentration of the indexed library, using AccuPrime Pfx polymerase (5µl of library template, 2 units AccuPrime Pfx DNA polymerase by Invitrogen, 1 unit of readymade 10x PCR mastermix, and 0,3µM of primers IS5 and IS6, for each 100µl reaction) with thermal profile of 2 min denaturation at 95°C, 3 to 9 cycles consisting of 15 sec denaturation at 95°C, 30 sec annealing at 60°C, 2 min elongation at 68°C and 5 min elongation at 68°C. The amplified libraries were purified using MinElute spin columns with the standard protocol provided by the manufacturer (Qiagen), and quantified for sequencing using an Agilent 2100 Bioanalyzer DNA 1000 chip.

For the modern Saami individual, total DNA was phenol-chloroform extracted and physically sheared using COVARIS fragmentation. A modified Illumina library preparation was performed using blunt-end repair followed by A-tailing of the 3’-end and ligation of forked adapters. Indexing PCR was followed by excision of fragments ranging from 500-600bp from a 2% agarose gel.

### Capture & Sequencing

We used the in-solution capture procedure from ref.^4^ to enrich our libraries for reads overlapping with 1,237,207 variable positions in the human genome. The captured libraries were sequenced (75bp single-end, plus additional paired end for the three non-UDG libraries of the Levänluhta individuals) on an Illumina HighSeq 4000 instrument at the Max Planck Institute for the Science of Human History in Jena. Out of the 13 originally processed Iron Age samples from Finland, seven proved to be of adequate quality to be used in downstream analyses. The modern Saami genome was sequenced in on a Genome Analyser II (8 lanes, 125bp paired-end) at the Max Planck Institute for Evolutionary Anthropology in Leipzig.

### Processing of sequenced reads

We used EAGER^56^ to process the sequenced reads, using the default parameters (see below) for human, UDG-half treated, single-end data, when processing the UDG-half libraries for all individuals. Specifically, AdapterRemoval was used to trim the sequencing adapters from our reads, with a minimum overlap of 1 bp, and using a minimum base quality of 20 and minimum sequence length of 30 bp. BWA aln (version 0.7.12-r1039) was used to map the reads to the hs37d5 human reference sequence, with a seed length (-l) of 32, max number of differences (-n) of 0.01 while doing no quality filtering. Duplicate removal was carried out using DeDup v0.12.1. Terminal base deamination damage calculation was done using mapDamage, specifying a length (-l) of 100 bp (Supplementary Table 1). For downstream analyses, we used bamutils (version 1.0.13) TrimBam to trim two bases at the start and end of all reads. This procedure eliminates the positions that are affected by deamination, thus removing genotyping errors that could arise due to ancient DNA damage.

For the untreated libraries of the three Levänluhta individuals, two rounds of sequencing were carried out, which were processed using EAGER with the above parameters, but specifying no UDG treatment and the correct sequencing type between the libraries. The merged reads were extracted from the resulting bam files, and merged with the bam file containing reads from the single end sequence run using samtools merge (version 1.3).

The data used to estimate mtDNA contamination for the three Levänluhta individuals were processed using EAGER: Adapter Removal (minimum overlap of 1 bp, minimum base quality of 20, minimum sequence length of 30, and with the option to “Keep only merged reads” activated) was used to merge reads (for the PE sequence run) and trim adapters. CircularMapper^56^ was used to map to the RSRS^57^, and the resulting bams were merged using samtools (version 1.3).

The modern Sami genome was using Ibis and an in-house adapter trimming script. The resulting reads were then aligned to the hs37d5 human reference genome using bwa 0.5.9-r16 (parameters -e 20 -o 2 -n 0.01).

### Genotyping

We used a custom script to genotype the 15 ancient individuals. For each individual and each SNP on the 1240K panel, one read covering the SNP was drawn at random, and a pseudo-haploid call was made, i.e., the ancient individual was assumed homozygous for the allele on the randomly drawn read for the SNP in question. The script is available at https://github.com/stschiff/sequenceTools.git.

For the three Levänluhta libraries that did not undergo UDG treatment, we only genotyped transversions, thus eliminating artefacts of post-mortem damage from further analyses.

The shotgun genome of the modern Saami individual was genotyped using GATK (version 1.3-25-g32cdef9) Unified Genotyper after indel realignment. The variant calls were filtered for variants with a quality score above 30, and a custom script was used to convert the variants into EigenStrat format.

The data were merged with a large dataset consisting of 3844 ancient and modern individuals genotyped on the Human Origins and/or 1240K SNP arrays, using mergeit.

### Sex determination

To determine the genetic sex of each ancient individual we calculated the coverage on the autosomes as well as on each sex chromosome. We then calculated the ratio of X-and Y-coverage over the autosomal coverage and plotted the results (Supplementary Figure 1). Females are expected to have an x-rate of 1 and a y-rate of 0, while males are expected to have both x-and y-rate of 0.5^45^.

### Authentication

We performed a number of different tests to assure the ancient authenticity of our data. For male individuals, we investigated polymorphisms on the X chromosome^19^ using the ANGSD software package (version 0.910)^58^. This revealed robust contamination estimates for 2 male Bolshoy individuals, and 1 male Chalmny-Varre individual. All of these are below 1.6% contamination (Table 1). For the female individuals from these two sites, we note that they are projected close to the males in PCA space (Figure 2a, Supplementary Figure 3), suggesting limited effects of potential contamination. In addition, we generated a PMD-filtered data set^21^ for all individuals using pmdtools (v 0.60). We set a pmd-threshold of 3, which, according to the original publication^21^, effectively eliminates potential modern contaminants based on the absence of base modifications consistent with deamination. We then tested whether the filtered and non-filtered data sets form a clade using F_4_ statistics of the form *f*_4_(All reads, PMD-filtered reads; *X*, Chimp) (Supplementary Table 2), while substituting *X* for Atayal, French, Kalash, Karitiana, Mbuti, Papuan and Yoruba. These statistics yielded no significant results, consistent with no substantial contamination in the data.

To provide a more quantitative estimate of possible contamination in females, we used ContamMix (version 1.0-10)^59^ to estimate mitochondrial contamination. We extracted the reads mapping to the mitochondrial reference for each of the ancient individuals using samtools (version 1.3). We then generated a mitochondrial consensus sequence for each of the ancient individuals using Geneious (version 10.0.9), and calling “N” for all sites with coverage lower than 5. Finally, all mitochondrial reads were aligned to their respective consensus sequence, using bwa aln (version 0.7.12-r1039) with a maximum number of differences in the seed (-k) set to 5 and the maximum number of differences (-n) to 10, and bwa samse (version 0.7.12-r1039). A multiple alignment of the consensus sequence and a reference set of 311 mitochondrial genomes^60^ was generated, using mafft (version v7.305b) with the “--auto” parameter. The read alignment, as well as the multiple alignment of the consensus and the 311 reference mitochondrial genomes were then provided to ContamMix. We report the mode a posteriori of contamination, along with the upper and lower bounds of the 95% posterior interval (Table 1).

For additional authentication, we ran supervised ADMIXTURE^20^ (version 1.3.0) for all samples using the six present-day populations above as defined genetic clusters, to look for large differences in genetic clustering among individuals from the same site (Supplementary Figure 2). We do not see significant differences (within our resolution) in the ancestry patterns between ancient individuals from the same site.

Finally, we projected PMD-filtered and non-filtered datasets on the same set of principal components constructed on modern European populations, using smartpca, to ensure that individuals are projected in close proximity regardless of PMD-filtering. This was possible for all samples with UDG-half treatment, except for the individuals from Levänluhta, which have too low damage to apply PMD-filtering. We therefore relied on the non-UDG libraries (using transversions only) that were generated for three individuals from the same site and used in the main analysis. We found that within expected noise due to a low number of SNPs, all samples show consistency between the filtered and non-filtered data sets, suggesting that contamination in all of these samples is low (Supplementary Figure 3a&b). Four additional individuals from Levänluhta were excluded from the main analysis and from this authentication test because of low coverage (<15,000 covered SNPs) and lack of non-UDG libraries.

### Principal component analysis

We used smartpca (version #13050) to carry out Principal Component (PC) analysis, using the “lsqproject: YES” parameter.

For the Eurasian PCA (Figure 2a), the following populations were used to construct principal components: Abkhasian, Adygei, Albanian, Altaian, Ami, Armenian, Atayal, Avar.SG, Azeri_WGA, Balkar, Balochi, Basque, BedouinA, BedouinB, Belarusian, Borneo, Brahui, Bulgarian, Buryat.SG, Cambodian, Canary_Islanders, Chechen, Chuvash, Croatian, Cypriot, Czech, Dai, Daur, Dolgan, Druze, English, Estonian, Even, Finnish, French, Georgian, Greek, Han, Hazara, Hezhen, Hungarian, Icelandic, Iranian, Italian_North, Italian_South, Japanese, Jew_Ashkenazi, Jew_Georgian, Jew_Iranian, Jew_Iraqi, Jew_Libyan, Jew_Moroccan, Jew_Tunisian, Jew_Turkish, Jew_Yemenite, Jordanian, Kalash, Kalmyk, Kinh, Korean, Kumyk, Kurd_WGA, Kyrgyz, Lahu, Lebanese, Lezgin, Lithuanian, Makrani, Mala, Maltese, Mansi, Miao, Mongola, Mordovian, Naxi, Nganasan, Nogai, North_Ossetian.DG, Norwegian, Orcadian, Oroqen, Palestinian, Pathan, Russian, Saami.DG, Saami_WGA, Sardinian, Saudi, Scottish, Selkup, Semende, She, Sherpa.DG, Sicilian, Spanish, Spanish_North, Syrian, Tajik, Thai, Tibetan.DG, Tu, Tubalar, Tujia, Turkish, Turkmen, Tuvinian, Ukrainian, Ulchi, Uygur, Uzbek, Xibo, Yakut, Yi, Yukagir.

For the European PCA (Supplementary Figure 3a &b), the following populations were used to construct principal components: Abkhasian, Adygei, Albanian, Armenian, Assyrian, Balkar, Basque, BedouinA, BedouinB, Belarusian, Bulgarian, Canary_Islander, Chechen, Chuvash, Croatian, Cypriot, Czech, Druze, English, Estonian, Finnish, French, Georgian, German, Greek, Hungarian, Icelandic, Iranian, Irish, Irish_Ulster, Italian_North, Italian_South, Jew_Ashkenazi, Jew_Georgian, Jew_Iranian, Jew_Iraqi, Jew_Libyan, Jew_Moroccan, Jew_Tunisian, Jew_Turkish, Jew_Yemenite, Jordanian, Kumyk, Lebanese, Lebanese_Christian, Lebanese_Muslim, Lezgin, Lithuanian, Maltese, Mordovian, North_Ossetian, Norwegian, Orcadian, Palestinian, Polish, Romanian, Russian, Sardinian, Saudi, Scottish, Shetlandic, Sicilian, Sorb, Spanish, Spanish_North, Syrian, Turkish, Ukrainian.

### ADMIXTURE analysis

ADMIXTURE was run with version 1.3.0, following LD pruning using plink (v1.90b3.29) with a window size of 200, a step size of 5 and an R^2^ threshold of 0.5. The populations used were: Albanian, Anatolia_Neolithic, Basque, Belarusian, BOO001, BOO002, BOO003, BOO004, BOO005, BOO006, Bulgarian, Chuvash, CHV001, CHV002, Croatian, Cypriot, Czech, EHG (Karelia + Samara), English, Estonian, Finnish, French, Greek, Hungarian, Icelandic, Iran_Chalcolithic, Italian_North, Italian_South, JK1963, JK1967, JK1968, JK1970, JK2065, JK2066, JK2067, Lithuanian, Loschbour_published.DG, MA1_HG.SG, ModernSaami, Mordovian, Nganasan, Norwegian, Poltavka, PPNB, PPNC, Russian, Saami.DG, Samara_Eneolithic, Sardinian, Scottish, SHG, Spanish, Spanish_North, Srubnaya, Stuttgart_published.DG, Ukrainian, WHG, Yamnaya_Samara.

While we technically find that K=2 results in the lowest Cross-Validation error, K=3 produced a similarly low error, and provided information more consistent with both previous publications^1^ and other analyses performed here (PCA, F statistiscs, qpAdm). In particular, at K=2 we already see the Siberian component shown in purple in Figure 2b, but no structure within European populations is observed at K=2. We therefore show K=3 as a reasonable descriptive analysis of our dataset.

### Y chromosome haplotyping

We assigned ancient males to Y haplogroups using the yHaplo program^61^. In short, this program provides an automated search through the Y haplogroup tree (as provided within yHaplo, as accessed from ISOGG on 04 Jan 2016) from the root to the downstream branch based on the presence of derived alleles and assign the most downstream haplogroup with derived alleles. For about 15,000 Y-chromosomal SNPs present both in our capture panel and in two published data sets^62,63^, we randomly sampled a single base and used it as a haploid genotype. We used a custom script to convert EigenStrat genotypes to the yHaplo format. We report the most downstream haplogroup provided by the program (Table 1). We also manually checked derived status and missingness in mutations defining the designated haplogroup because missing information might lead to a premature stop in its automated search.

### Mitochondrial haplotyping

We imported the trimmed mitochondrial reads for each individual with mapping quality > 30 into Geneious (v. 10.0.9) and reassembled those reads to the RSRS^57^, using the Geneious mapper, with medium sensitivity and 5 iterations. We used the in-built automated variant caller within Geneious to find mitochondrial polymorphisms with a minimum coverage of 3 and a minimum Variant Frequency of 0.67. The resulting variants were exported to Excel and manually compared to the SNPs reported in the online mtDNA phylogeny (mtDNA tree Build 17, 18 Feb 2016, http://www.phylotree.org/). Nucleotide positions 309.1C(C), 315.1C, AC indels at 515-522, 16182C, 16183C, 16193.1C(C) and 16519 were masked and not included in our haplotype calls.

### Phenotypic SNPs

We used samtools (v. 1.3) mpileup, filtering for map-(-Q) and base-(-q) quality of 30, deactivating per-Base Alignment Quality (-B), on the trimmed bam files, to generate a pileup of reads mapping to a set of 43 Phenotypic SNPs^4,32,33,64^ that are part of our genome capture panel. A custom script was used to parse the pileup into a table containing the number of reads supporting each allele (Supplementary Table 7).

## Acknowledgements

We thank everyone who contributed to the archaeological excavations, Cosimo Posth for carrying out laboratory work, the lab technicians involved in this project, and the sequencing team at the Max Planck Institute for the Science of Human History. We would like to also thank the sequencing team at Max Planck Institute of Evolutionary Anthropology for the sequencing of the modern Saami genome. This project was funded by Emil Aaltonen Foundation, Jane and Aatos Erkko Foundation, the Kone Foundation, and the Max Planck Society.

## Author Contributions

S.S., J.Kr., and W.H. supervised the study. A.Wes., V.M., V.K., A.S., P.O., O.B., W.H. and J.Kr. assembled the collection of archaeological samples. A.S. and S.P. collected the modern Saami sample. K.M. performed laboratory work. K.M. and T.C.L. supervised ancient DNA sequencing and post-sequencing bioinformatics for the ancient individuals. A.Wei. performed the laboratory work of the modern Saami genome. M.O. carried out the processing of the sequenced reads and generating the genotypes of the modern Saami genome. J.Ke. and S.P. supervised sequencing of the modern Saami and post-sequencing bioinformatics. T.C.L., K.M., E.S., C.J. and S.S. analysed genetic data. T.C.L., K.M., E.S., W.H., J.Kr. and S.S. wrote the manuscript with additional input from all other co-authors.

## Author information

Data from all ancient individuals and from the modern Saami individual is available under accession numbers XXX. The authors declare no competing financial interests.

