## Supplementary Materials for "Ancient Fennoscandian genomes reveal origin and spread of Siberian ancestry in Europe"

### Supplementary Material for Ancient Fennoscandian genomes reveal origin and spread of Siberian ancestry in Europe

#### Supplementary Figures

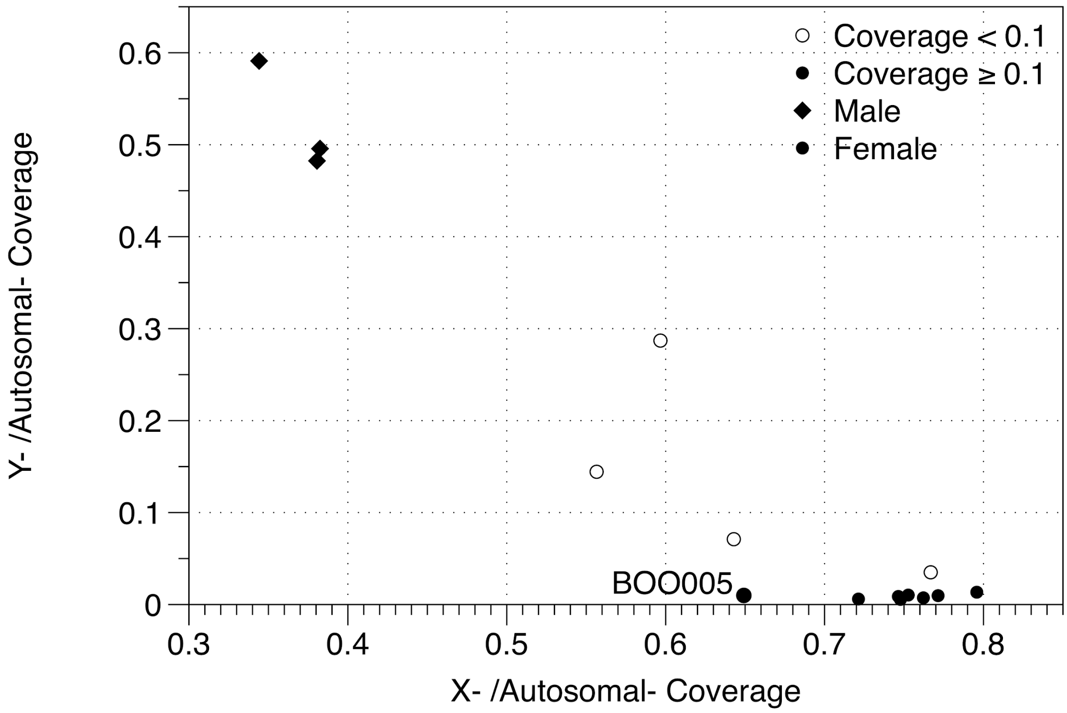


Supplementary Figure 1. Sex Determination. X-chromosomal coverage vs. Y-chromosomal coverage, normalised by autosomal coverage. Low autosomal coverage individuals (n = 4, excluded from downstream analyses) are shown as empty circles, and higher coverage individuals (n = 11) as filled circles. BOO005 is highlighted because of its low relative X coverage.

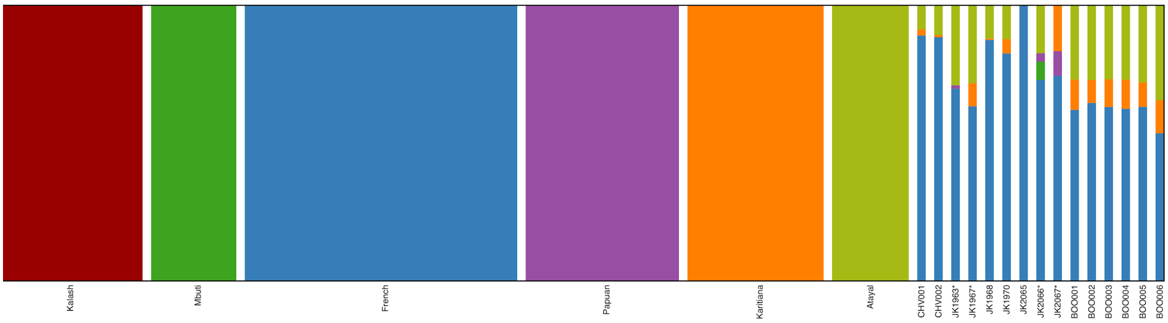


Supplementary Figure 2. Supervised ADMIXTURE. Individuals excluded from further analyses due to low coverage are signified with an asterisk. Among higher coverage genomes, the results within each population are homogeneous, with the exception of the outlier JK2065 within Levänluhta.

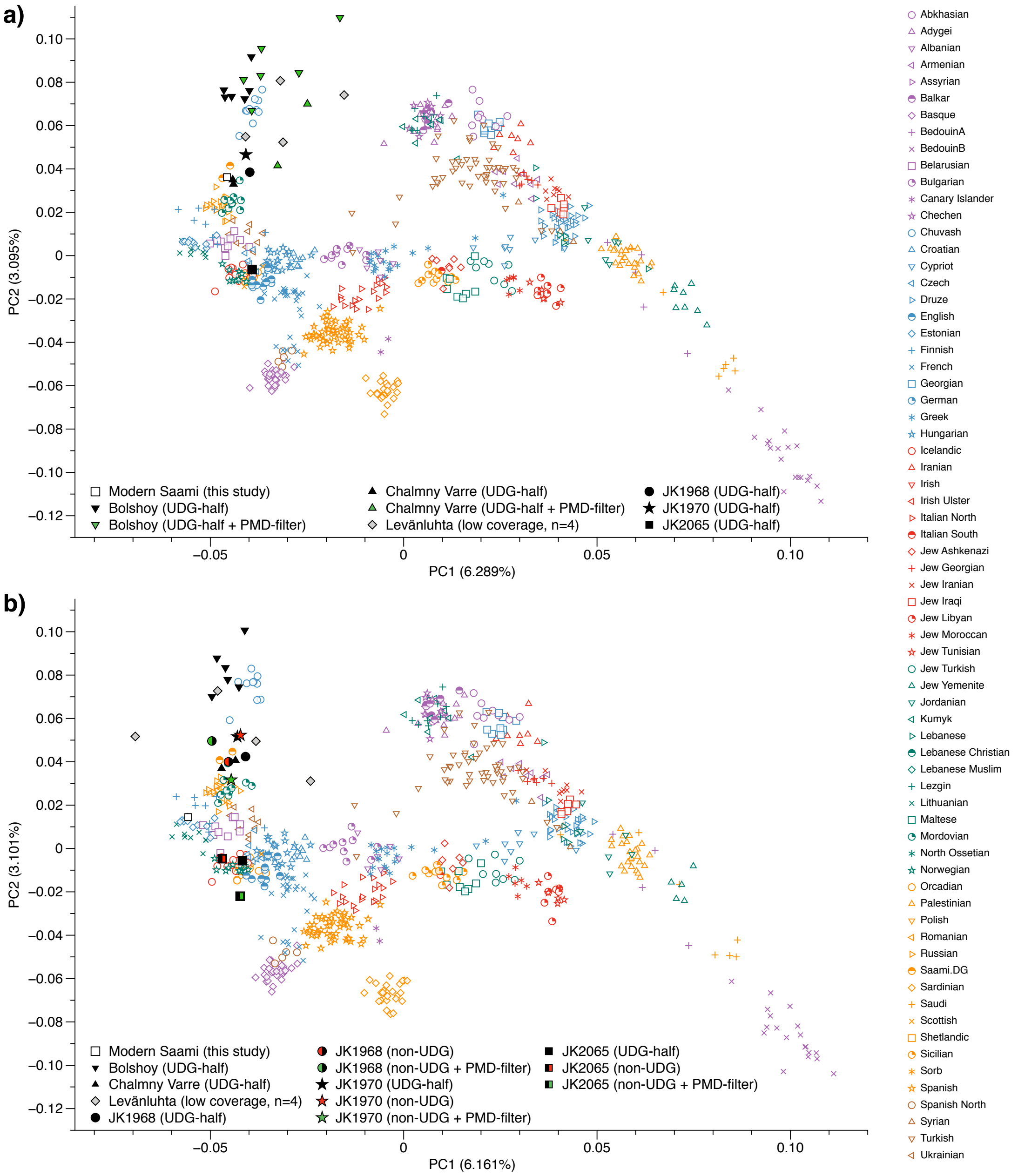


Supplementary Figure 3. **a)** PCA plot of Europe with individuals from this study projected on principal components constructed on the populations in the legend. For each individual from Bolshoy and Chalmny Varre, an additional projection is shown for the PMD-filtered dataset. Ancient individuals with fewer than 15,000 covered SNPs shown in grey. b) PCA plot of Europe with individuals from this study projected on principal components constructed using only transversion SNPs of the populations in the legend. For the three Levänluhta individuals with more than 15,000 covered SNPs, additional point for the PMD-filtered and the non-filtered datasets of the non-UDG treated libraries are shown. Ancient individuals with fewer than 15,000 covered SNPs shown in grey.

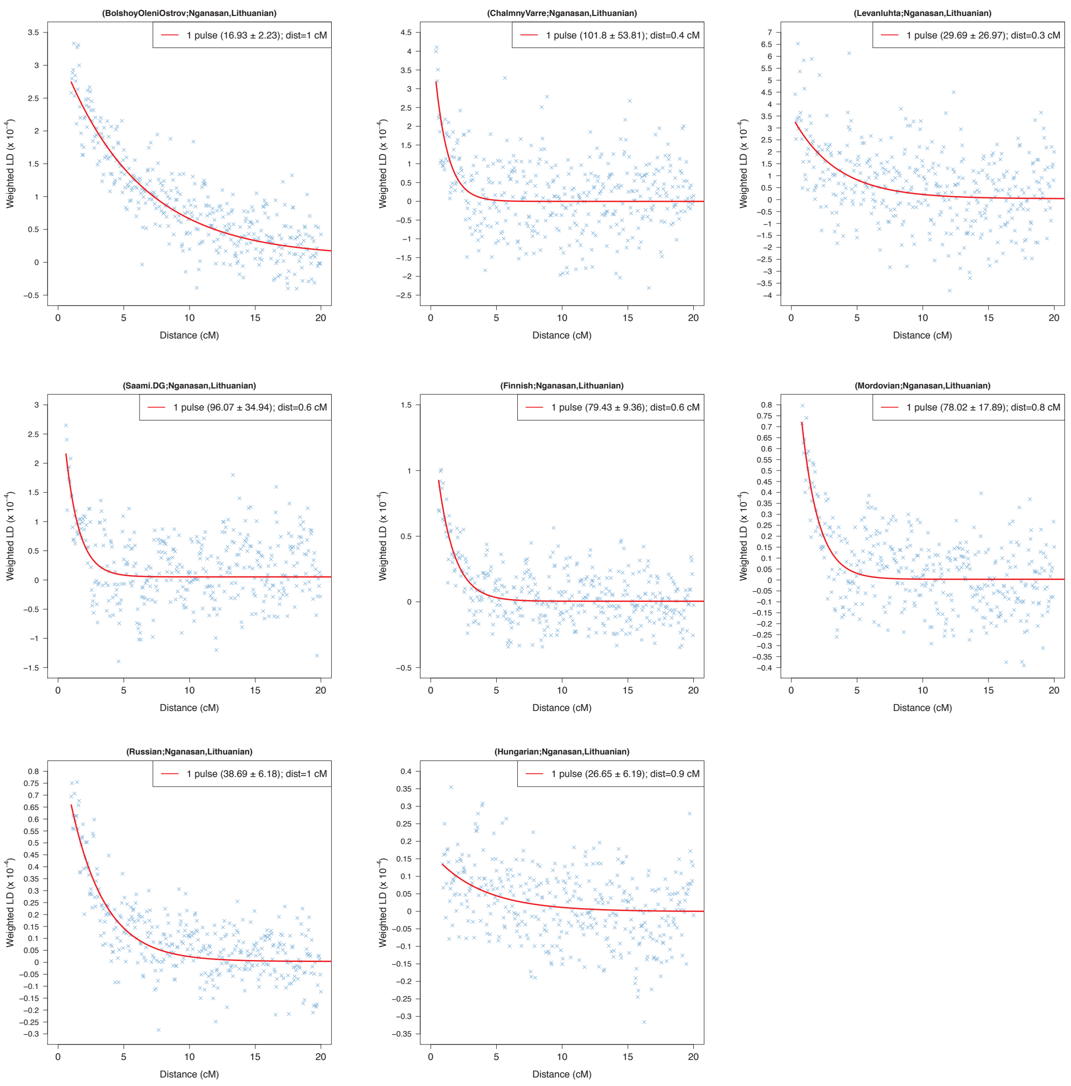


Supplementary Figure 4. Linkage Disequilibrium decay curves for ancient and modern populations. Minimum distance in cM used was the lowest distance for which ALDER provided results.

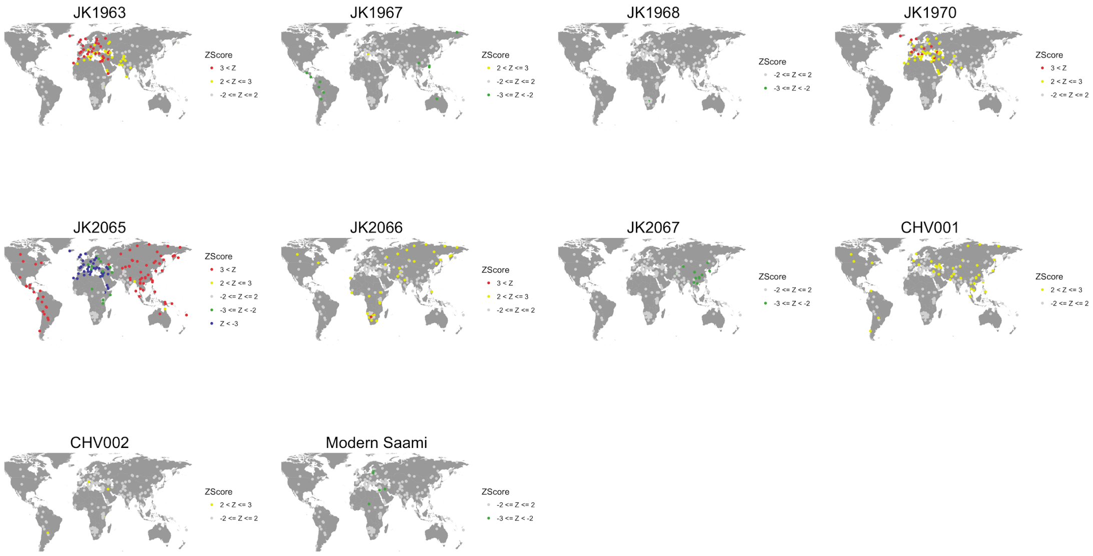


Supplementary Figure 5. $f_4$(Saami, Test; X, Mbuti) for multiple worldwide populations *X*, using ancient and modern individuals from this study as *Test*. Points are coloured based on bins of Z Score values, with warmer colours indicating higher affinity in Saami than in the *Test*, and vice versa for colder colours. Grey points indicate similar affinities toward population *X*.

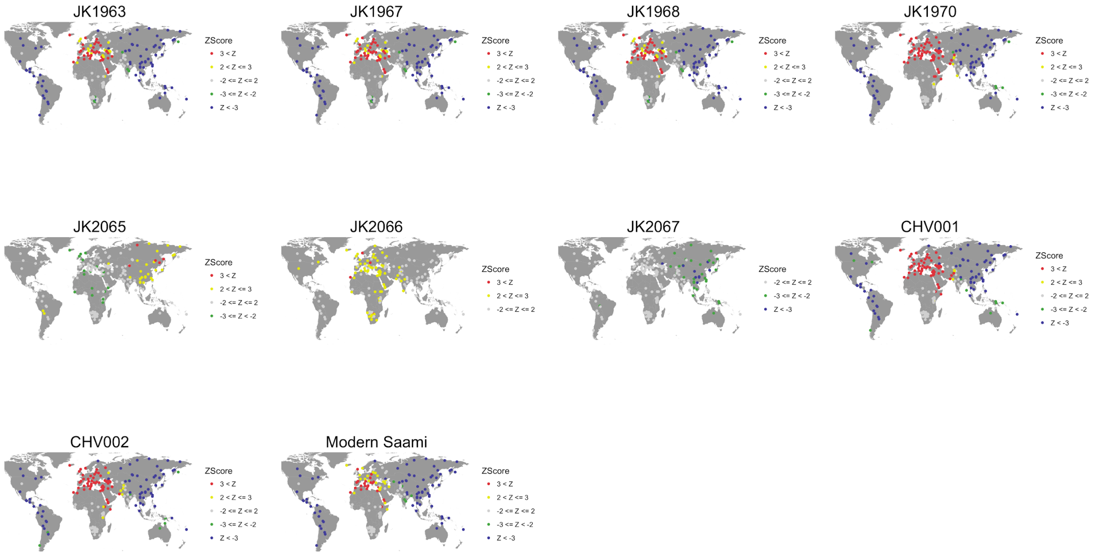


Supplementary Figure 6. $f_4$(Finnish, Test; X, Mbuti) for multiple worldwide populations *X*, using ancient and modern individuals from this study as *Test*. Points are coloured based on bins of Z Score values, with warmer colours indicating higher affinity in Finns than in the *Test*, and vice versa for colder colours. Grey points indicate similar affinities toward population *X*.

#### Supplementary Tables

Supplementary Table 1. Deamination damage per library.

| Library ID | UDG treatment | Damage 5' 1st base | Damage 5' 2nd base | Damage 3' 2nd base | Damage 3' 1st base |
|---|---|---|---|---|---|
| CHV001 | Half | 0.0287 | 0.0074 | 0.0121 | 0.031 |
| CHV002 | Half | 0.0189 | 0.0062 | 0.0111 | 0.0244 |
| JK1963 | Half | 0.0259 | 0.0071 | 0.0111 | 0.0294 |
| JK1967 | Half | 0.0104 | 0.0048 | 0.009 | 0.0151 |
| JK1968 | Half | 0.0284 | 0.0054 | 0.0088 | 0.0282 |
| JK1968.nonUDG | None | 0.094 | 0.0575 | 0.0521 | 0.0865 |
| JK1970 | Half | 0.0239 | 0.0069 | 0.0109 | 0.029 |
| JK1970.nonUDG | None | 0.0953 | 0.0745 | 0.073 | 0.0937 |
| JK2065 | Half | 0.0123 | 0.0045 | 0.0069 | 0.0155 |
| JK2065.nonUDG | None | 0.0471 | 0.0452 | 0.0448 | 0.0487 |
| JK2066 | Half | 0.0146 | 0.0041 | 0.0059 | 0.0171 |
| JK2067 | Half | 0.0104 | 0.004 | 0.0065 | 0.0129 |
| BOO001 | Half | 0.0283 | 0.0075 | 0.0107 | 0.0302 |
| BOO002 | Half | 0.0402 | 0.0078 | 0.0113 | 0.0333 |
| BOO003 | Half | 0.0411 | 0.0073 | 0.0108 | 0.0398 |
| BOO004 | Half | 0.0382 | 0.0085 | 0.0117 | 0.0344 |
| BOO005 | Half | 0.045 | 0.0066 | 0.0108 | 0.0421 |
| BOO006 | Half | 0.0381 | 0.0063 | 0.0094 | 0.0302 |

Supplementary Table 2. $f_4$(All reads, PMD-filtered reads; X, Chimp).

| Ancient Individual | Population X | $f_4$ | Z Score | # of SNPs |
|---|---|---|---|---|
| BOO001 | Mbuti | -0.000699 | -0.683 | 14602 |
| BOO001 | Kalash | 0.000056 | 0.048 | 14602 |
| BOO001 | French | -0.000361 | -0.315 | 14602 |
| BOO001 | Papuan | 0.000255 | 0.219 | 14602 |
| BOO001 | Karitiana | -0.000202 | -0.157 | 14602 |
| BOO001 | Atayal | -0.000208 | -0.169 | 14602 |
| BOO001 | Yoruba | 0.000191 | 0.187 | 14602 |
| BOO002 | Mbuti | -0.000019 | -0.018 | 10659 |
| BOO002 | Kalash | 0.000809 | 0.689 | 10659 |
| BOO002 | French | 0.00083 | 0.691 | 10659 |
| BOO002 | Papuan | 0.000766 | 0.608 | 10659 |
| BOO002 | Karitiana | 0.000139 | 0.104 | 10659 |
| BOO002 | Atayal | 0.000433 | 0.361 | 10659 |
| BOO002 | Yoruba | 0.000433 | 0.431 | 10659 |
| BOO003 | Mbuti | 0.000548 | 0.609 | 15523 |
| BOO003 | Kalash | 0.000139 | 0.137 | 15523 |
| BOO003 | French | -0.000264 | -0.268 | 15523 |
| BOO003 | Papuan | 0.000085 | 0.081 | 15523 |
| BOO003 | Karitiana | -0.000508 | -0.457 | 15523 |
| BOO003 | Atayal | -0.000454 | -0.409 | 15523 |
| BOO003 | Yoruba | -0.000085 | -0.095 | 15523 |
| BOO004 | Mbuti | 0.000472 | 0.687 | 39822 |
| BOO004 | Kalash | 0.000033 | 0.041 | 39822 |
| BOO004 | French | 0.000401 | 0.514 | 39822 |
| BOO004 | Papuan | 0.000471 | 0.592 | 39822 |
| BOO004 | Karitiana | 0.000613 | 0.72 | 39822 |
| BOO004 | Atayal | 0.000085 | 0.105 | 39822 |
| BOO004 | Yoruba | 0.000053 | 0.077 | 39822 |
| BOO005 | Mbuti | 0.00051 | 0.826 | 47172 |
| BOO005 | Kalash | 0.000485 | 0.685 | 47172 |
| BOO005 | French | 0.000657 | 0.948 | 47172 |
| BOO005 | Papuan | 0.000404 | 0.557 | 47172 |
| BOO005 | Karitiana | 0.000164 | 0.207 | 47172 |
| BOO005 | Atayal | 0.000516 | 0.683 | 47172 |
| BOO005 | Yoruba | 0.000812 | 1.3 | 47172 |
| BOO006 | Mbuti | -0.000676 | -0.717 | 13388 |
| BOO006 | Kalash | 0.000836 | 0.785 | 13388 |
| BOO006 | French | 0.000625 | 0.613 | 13388 |
| BOO006 | Papuan | 0.000174 | 0.161 | 13388 |
| BOO006 | Karitiana | 0.000343 | 0.287 | 13388 |
| BOO006 | Atayal | 0.000492 | 0.454 | 13388 |
| BOO006 | Yoruba | -0.000467 | -0.51 | 13388 |
| CHV001 | Mbuti | -0.000999 | -1.057 | 17553 |
| CHV001 | Kalash | -0.00077 | -0.748 | 17553 |

| | | | | |
|---|---|---|---|---|
| CHV001 | French | -0.000459 | -0.462 | 17553 |
| CHV001 | Papuan | -0.000689 | -0.626 | 17553 |
| CHV001 | Karitiana | 0.000185 | 0.166 | 17553 |
| CHV001 | Atayal | -0.000835 | -0.773 | 17553 |
| CHV001 | Yoruba | -0.000724 | -0.79 | 17553 |
| CHV002 | Mbuti | 0.000267 | 0.227 | 5082 |
| CHV002 | Kalash | 0.000048 | 0.037 | 5082 |
| CHV002 | French | -0.000152 | -0.117 | 5082 |
| CHV002 | Papuan | -0.00026 | -0.193 | 5082 |
| CHV002 | Karitiana | 0.000543 | 0.388 | 5082 |
| CHV002 | Atayal | -0.000041 | -0.029 | 5082 |
| CHV002 | Yoruba | -0.000483 | -0.415 | 5082 |
| JK1963 | Mbuti | 0 | 0 | 0 |
| JK1963 | Kalash | 0 | 0 | 0 |
| JK1963 | French | 0 | 0 | 0 |
| JK1963 | Papuan | 0 | 0 | 0 |
| JK1963 | Karitiana | 0 | 0 | 0 |
| JK1963 | Atayal | 0 | 0 | 0 |
| JK1963 | Yoruba | 0 | 0 | 0 |
| JK1967 | Mbuti | 0 | 0 | 0 |
| JK1967 | Kalash | 0 | 0 | 0 |
| JK1967 | French | 0 | 0 | 0 |
| JK1967 | Papuan | 0 | 0 | 0 |
| JK1967 | Karitiana | 0 | 0 | 0 |
| JK1967 | Atayal | 0 | 0 | 0 |
| JK1967 | Yoruba | 0 | 0 | 0 |
| JK1968 | Mbuti | 0.002381 | 1.978 | 5729 |
| JK1968 | Kalash | 0.001631 | 1.247 | 5729 |
| JK1968 | French | 0.001069 | 0.85 | 5729 |
| JK1968 | Papuan | 0.002575 | 1.954 | 5729 |
| JK1968 | Karitiana | 0.001855 | 1.326 | 5729 |
| JK1968 | Atayal | 0.001924 | 1.359 | 5729 |
| JK1968 | Yoruba | 0.001679 | 1.415 | 5729 |
| JK1970 | Mbuti | 0.001332 | 1.202 | 5788 |
| JK1970 | Kalash | 0.000674 | 0.554 | 5788 |
| JK1970 | French | 0.000424 | 0.36 | 5788 |
| JK1970 | Papuan | -0.000314 | -0.25 | 5788 |
| JK1970 | Karitiana | 0.000754 | 0.599 | 5788 |
| JK1970 | Atayal | 0.00069 | 0.556 | 5788 |
| JK1970 | Yoruba | 0.001119 | 1.047 | 5788 |
| JK2065 | Mbuti | 0.001841 | 1.029 | 2257 |
| JK2065 | Kalash | 0.001897 | 1.025 | 2257 |
| JK2065 | French | 0.002606 | 1.347 | 2257 |
| JK2065 | Papuan | 0.001431 | 0.701 | 2257 |
| JK2065 | Karitiana | 0.003007 | 1.44 | 2257 |
| JK2065 | Atayal | 0.001972 | 0.898 | 2257 |
| JK2065 | Yoruba | 0.001445 | 0.826 | 2257 |
| JK2066 | Mbuti | 0 | 0 | 0 |
| JK2066 | Kalash | 0 | 0 | 0 |
| JK2066 | French | 0 | 0 | 0 |
| JK2066 | Papuan | 0 | 0 | 0 |
| JK2066 | Karitiana | 0 | 0 | 0 |
| JK2066 | Atayal | 0 | 0 | 0 |
| JK2066 | Yoruba | 0 | 0 | 0 |
| JK2067 | Mbuti | 0 | 0 | 0 |
| JK2067 | Kalash | 0 | 0 | 0 |
| JK2067 | French | 0 | 0 | 0 |
| JK2067 | Papuan | 0 | 0 | 0 |
| JK2067 | Karitiana | 0 | 0 | 0 |
| JK2067 | Atayal | 0 | 0 | 0 |

Supplementary Table 3. $f_3$(Test; Siberian source, European source). the $f_3$ with the most negative Z Score is highlighted in bold for each populations, where applicable.

| Test | Siberian | European | $f_3$ | Z Score | # SNPs |
|---|---|---|---|---|---|
| All Saami* | Nganasan | French | 0.003588 | 2.704 | 441155 |
| All Saami* | Nganasan | Icelandic | 0.002458 | 1.780 | 422594 |
| All Saami* | Nganasan | Lithuanian | 0.002818 | 2.081 | 419356 |
| All Saami* | Nganasan | Norwegian | 0.002898 | 2.114 | 422620 |
| All Saami* | Mansi | French | 0.015804 | 11.816 | 438903 |
| All Saami* | Mansi | Icelandic | 0.015392 | 11.307 | 421190 |
| All Saami* | Mansi | Lithuanian | 0.015589 | 11.649 | 418774 |
| All Saami* | Mansi | Norwegian | 0.015416 | 11.535 | 421289 |
| All Saami* | Bolshoy | French | 0.012524 | 9.419 | 401267 |
| All Saami* | Bolshoy | Icelandic | 0.012438 | 9.265 | 379480 |
| All Saami* | Bolshoy | Lithuanian | 0.013625 | 10.075 | 375339 |
| All Saami* | Bolshoy | Norwegian | 0.013379 | 9.852 | 379088 |
| Estonian | Nganasan | French | 0.000198 | 0.445 | 443041 |
| Estonian | Nganasan | Icelandic | -0.00032 | -0.631 | 429850 |
| **Estonian** | **Nganasan** | **Lithuanian** | **-0.001771** | **-3.425** | **427789** |
| Estonian | Nganasan | Norwegian | 0.000359 | 0.722 | 429940 |
| Estonian | Mansi | French | 0.000452 | 1.257 | 440774 |
| Estonian | Mansi | Icelandic | 0.000629 | 1.617 | 427699 |
| Estonian | Mansi | Lithuanian | -0.000979 | -2.419 | 425886 |
| Estonian | Mansi | Norwegian | 0.000905 | 2.351 | 427790 |
| Estonian | Bolshoy | French | -0.001054 | -2.607 | 403038 |
| Estonian | Bolshoy | Icelandic | -0.000519 | -1.164 | 387741 |
| Estonian | Bolshoy | Lithuanian | -0.00117 | -2.520 | 385139 |
| Estonian | Bolshoy | Norwegian | 0.000614 | 1.338 | 387751 |
| Finnish | Nganasan | French | -0.004643 | -9.166 | 442567 |
| **Finnish** | **Nganasan** | **Icelandic** | **-0.005401** | **-9.621** | **427954** |
| Finnish | Nganasan | Lithuanian | -0.005166 | -8.812 | 426231 |
| Finnish | Nganasan | Norwegian | -0.004848 | -8.574 | 428161 |
| Finnish | Mansi | French | -0.001386 | -3.424 | 440277 |
| Finnish | Mansi | Icelandic | -0.001453 | -3.253 | 426028 |
| Finnish | Mansi | Lithuanian | -0.001374 | -2.959 | 424645 |
| Finnish | Mansi | Norwegian | -0.001301 | -2.840 | 426248 |
| Finnish | Bolshoy | French | -0.002922 | -6.654 | 402958 |
| Finnish | Bolshoy | Icelandic | -0.002698 | -5.581 | 386418 |
| Finnish | Bolshoy | Lithuanian | -0.001631 | -3.066 | 384134 |
| Finnish | Bolshoy | Norwegian | -0.001661 | -3.333 | 386203 |
| **Hungarian** | **Nganasan** | **French** | **-0.001262** | **-4.169** | **446371** |
| Hungarian | Nganasan | Icelandic | -0.000749 | -1.812 | 437464 |
| Hungarian | Nganasan | Lithuanian | -0.000436 | -1.020 | 436905 |
| Hungarian | Nganasan | Norwegian | -0.000318 | -0.785 | 437888 |
| Hungarian | Mansi | French | -0.000834 | -3.794 | 444139 |
| Hungarian | Mansi | Icelandic | 0.000367 | 1.228 | 435158 |
| Hungarian | Mansi | Lithuanian | 0.000524 | 1.671 | 434697 |
| Hungarian | Mansi | Norwegian | 0.000399 | 1.393 | 435533 |
| Hungarian | Bolshoy | French | -0.001262 | -4.677 | 406930 |
| Hungarian | Bolshoy | Icelandic | 0.000269 | 0.757 | 397199 |
| Hungarian | Bolshoy | Lithuanian | 0.001365 | 3.496 | 396443 |
| Hungarian | Bolshoy | Norwegian | 0.001176 | 3.155 | 397522 |
| Modern Saami | Nganasan | French | 0.056315 | 8.011 | 440449 |
| Modern Saami | Nganasan | Icelandic | 0.055309 | 7.888 | 420374 |
| Modern Saami | Nganasan | Lithuanian | 0.054694 | 7.777 | 416406 |
| Modern Saami | Nganasan | Norwegian | 0.055971 | 7.970 | 420260 |
| Modern Saami | Mansi | French | 0.067798 | 9.408 | 438095 |
| Modern Saami | Mansi | Icelandic | 0.067585 | 9.407 | 419053 |
| Modern Saami | Mansi | Lithuanian | 0.066793 | 9.291 | 415996 |
| Modern Saami | Mansi | Norwegian | 0.067788 | 9.441 | 419031 |
| Modern Saami | Bolshoy | French | 0.064081 | 8.914 | 399764 |
| Modern Saami | Bolshoy | Icelandic | 0.064262 | 8.935 | 375157 |
| Modern Saami | Bolshoy | Lithuanian | 0.064566 | 8.951 | 369834 |
| Modern Saami | Bolshoy | Norwegian | 0.065519 | 9.112 | 374544 |
| Mordovian | Nganasan | French | -0.006169 | -13.753 | 443651 |
| Mordovian | Nganasan | Icelandic | -0.006308 | -12.383 | 430318 |
| **Mordovian** | **Nganasan** | **Lithuanian** | **-0.007452** | **-14.743** | **428647** |
| Mordovian | Nganasan | Norwegian | -0.00597 | -11.724 | 430787 |
| Mordovian | Mansi | French | -0.002921 | -8.312 | 441523 |
| Mordovian | Mansi | Icelandic | -0.002373 | -6.222 | 428477 |
| Mordovian | Mansi | Lithuanian | -0.003673 | -9.969 | 427165 |
| Mordovian | Mansi | Norwegian | -0.002434 | -6.339 | 428926 |

| | | | | | |
|---|---|---|---|---|---|
| Mordovian | Bolshoy | French | -0.003619 | -9.245 | 404494 |
| Mordovian | Bolshoy | Icelandic | -0.002769 | -6.277 | 389566 |
| Mordovian | Bolshoy | Lithuanian | -0.003101 | -6.753 | 387543 |
| Mordovian | Bolshoy | Norwegian | -0.001941 | -4.285 | 389938 |
| Russian | Nganasan | French | -0.00743 | -23.161 | 445788 |
| Russian | Nganasan | Icelandic | -0.007939 | -20.339 | 436162 |
| **Russian** | **Nganasan** | **Lithuanian** | **-0.008804** | **-21.719** | **435228** |
| Russian | Nganasan | Norwegian | -0.007252 | -18.830 | 436431 |
| Russian | Mansi | French | -0.003891 | -17.070 | 443954 |
| Russian | Mansi | Icelandic | -0.003713 | -13.190 | 434285 |
| Russian | Mansi | Lithuanian | -0.004733 | -16.274 | 433573 |
| Russian | Mansi | Norwegian | -0.003425 | -11.793 | 434584 |
| Russian | Bolshoy | French | -0.004841 | -17.610 | 407944 |
| Russian | Bolshoy | Icelandic | -0.004337 | -13.017 | 397811 |
| Russian | Bolshoy | Lithuanian | -0.004425 | -12.276 | 396731 |
| Russian | Bolshoy | Norwegian | -0.003199 | -9.000 | 398111 |
| Saami(SGDP) | Nganasan | French | 0.000914 | 0.510 | 434884 |
| **Saami(SGDP)** | **Nganasan** | **Icelandic** | **-0.000333** | **-0.182** | **415857** |
| Saami(SGDP) | Nganasan | Lithuanian | 0.000511 | 0.284 | 412432 |
| Saami(SGDP) | Nganasan | Norwegian | 0.000005 | 0.003 | 415843 |
| Saami(SGDP) | Mansi | French | 0.014073 | 7.908 | 432644 |
| Saami(SGDP) | Mansi | Icelandic | 0.013525 | 7.527 | 414586 |
| Saami(SGDP) | Mansi | Lithuanian | 0.014213 | 7.936 | 412029 |
| Saami(SGDP) | Mansi | Norwegian | 0.013453 | 7.636 | 414611 |
| Saami(SGDP) | Bolshoy | French | 0.011112 | 6.168 | 395613 |
| Saami(SGDP) | Bolshoy | Icelandic | 0.010859 | 6.033 | 373006 |
| Saami(SGDP) | Bolshoy | Lithuanian | 0.012541 | 6.859 | 368453 |
| Saami(SGDP) | Bolshoy | Norwegian | 0.011686 | 6.514 | 372515 |

*"All Saami" refers to a grouping including the 2 individuals from the SGDP (Saami(SGDP)) and the high-coverage modern Saami shotgun genome in this study (Modern Saami).

Supplementary Table 4. qpAdm models. Right populations used were Mbuti, Villabruna, Han, Onge, BedouinB and Mixe. Models with p-values above p=0.05 are highlighted in red.

| Test | p-value | WHG | Karelia HG | LBK EN | Yamnaya | Nganasan |
|---|---|---|---|---|---|---|
| BolshoyOleniOstrov | 1.71E-01 | 0.074 ±0.018 | 0.320 ±0.101 | -0.045 ±0.083 | 0.216 ±0.182 | 0.436 ±0.013 |
| ChalmnyVarre | 3.68E-01 | 0.169 ±0.023 | 0.159 ±0.126 | 0.266 ±0.109 | 0.158 ±0.234 | 0.248 ±0.018 |
| Levänluhta* | 1.85E-01 | 0.134 ±0.026 | 0.247 ±0.148 | 0.292 ±0.123 | 0.041 ±0.267 | 0.286 ±0.018 |
| CHV001 | 4.20E-01 | 0.171 ±0.027 | 0.182 ±0.141 | 0.264 ±0.124 | 0.145 ±0.266 | 0.238 ±0.021 |
| CHV002 | 2.27E-01 | 0.191 ±0.036 | -0.09 ±0.205 | 0.101 ±0.173 | 0.546 ±0.372 | 0.251 ±0.025 |
| Estonian | 1.04E-01 | 0.141 ±0.011 | 0.084 ±0.070 | 0.472 ±0.056 | 0.243 ±0.126 | 0.061 ±0.008 |
| Finnish | 3.70E-02 | 0.131 ±0.012 | 0.108 ±0.097 | 0.453 ±0.079 | 0.209 ±0.177 | 0.100 ±0.009 |
| French | 7.56E-02 | 0.084 ±0.009 | 0.057 ±0.052 | 0.665 ±0.043 | 0.159 ±0.095 | 0.035 ±0.006 |
| Hungarian | 1.29E-01 | 0.073 ±0.010 | 0.103 ±0.053 | 0.630 ±0.043 | 0.150 ±0.095 | 0.045 ±0.007 |
| JK1968 | 1.81E-01 | 0.096 ±0.041 | 0.472 ±0.186 | 0.487 ±0.160 | -0.311 ±0.332 | 0.255 ±0.027 |
| JK1970 | 2.22E-01 | 0.157 ±0.034 | 0.061 ±0.203 | 0.087 ±0.175 | 0.392 ±0.382 | 0.303 ±0.026 |
| JK2065 | 3.14E-01 | 0.177 ±0.044 | 0.591 ±0.267 | 0.834 ±0.254 | -0.664 ±0.561 | 0.063 ±0.045 |
| Lithuanian | 5.55E-02 | 0.147 ±0.013 | 0.218 ±0.090 | 0.586 ±0.072 | 0.006 ±0.163 | 0.042 ±0.010 |
| Mordovian | 1.00E-01 | 0.098 ±0.012 | 0.153 ±0.078 | 0.498 ±0.063 | 0.144 ±0.142 | 0.108 ±0.009 |
| Russian | 6.95E-02 | 0.105 ±0.009 | 0.093 ±0.067 | 0.438 ±0.052 | 0.252 ±0.120 | 0.112 ±0.007 |
| Saami.DG | 1.81E-01 | 0.101 ±0.026 | 0.468 ±0.134 | 0.563 ±0.113 | -0.424 ±0.243 | 0.292 ±0.019 |
| Italian_South | 8.20E-04 | -0.091 ±0.026 | -0.161 ±0.178 | 0.656 ±0.152 | 0.580 ±0.330 | 0.016 ±0.021 |
| Spanish | 8.36E-06 | 0.073 ±0.016 | -0.285 ±0.184 | 0.436 ±0.153 | 0.774 ±0.339 | 0.002 ±0.015 |
| CordedWare_Baltic | 1.32E-02 | 0.126 ±0.019 | -0.092 ±0.110 | 0.109 ±0.092 | 0.846 ±0.199 | 0.012 ±0.012 |
| Corded_Ware_Germany | 2.68E-01 | 0.068 ±0.016 | 0.249 ±0.082 | 0.422 ±0.070 | 0.243 ±0.152 | 0.018 ±0.011 |
| Kunda_LT | 8.37E-03 | 0.896 ±0.035 | 0.216 ±0.212 | 0.176 ±0.177 | -0.318 ±0.389 | 0.030 ±0.022 |
| Narva_LT | 1.12E-02 | 0.880 ±0.024 | 0.093 ±0.116 | 0.087 ±0.096 | -0.056 ±0.209 | -0.004 ±0.014 |
| BA_Baltic | 9.62E-02 | 0.262 ±0.014 | 0.078 ±0.074 | 0.308 ±0.062 | 0.346 ±0.134 | 0.007 ±0.009 |
| BOO001 | 4.65E-02 | 0.009 ±0.036 | 0.545 ±0.223 | 0.064 ±0.189 | -0.023 ±0.406 | 0.404 ±0.026 |
| BOO002 | 3.26E-02 | 0.087 ±0.035 | 0.368 ±0.200 | 0.128 ±0.175 | -0.011 ±0.371 | 0.429 ±0.027 |
| BOO003 | 7.70E-02 | 0.093 ±0.032 | 0.177 ±0.171 | -0.171 ±0.132 | 0.500 ±0.293 | 0.402 ±0.024 |
| BOO004 | 1.21E-01 | 0.061 ±0.031 | 0.236 ±0.194 | -0.072 ±0.160 | 0.364 ±0.347 | 0.411 ±0.024 |
| BOO005 | 1.43E-01 | 0.138 ±0.031 | 0.220 ±0.163 | -0.199 ±0.140 | 0.466 ±0.301 | 0.376 ±0.025 |
| BOO006 | 4.68E-02 | 0.077 ±0.031 | 0.303 ±0.217 | 0.006 ±0.176 | 0.070 ±0.385 | 0.544 ±0.024 |

*Levänluhta here refers to JK1968 & JK1970 grouped as one population.

Supplementary Table 5. $f_4$ (Finnish, Test; X, Mbuti) for multiple worldwide populations X. Z Scores highlighted by significance bin. Red: Z>3, yellow: 3≥Z>2, gray: 2≥Z≥-2, green: -2>Z≥-3 and blue: Z<-3. (Found in accompanying Excel spreadsheet.)

Supplementary Table 6. $f_4$ (Saami (SGDP), Test; X, Mbuti) for multiple worldwide populations X. Z Scores highlighted by significance bin. Red: Z>3, yellow: 3≥Z>2, gray: 2≥Z≥-2, green: -2>Z≥-3 and blue: Z<-3. (Found in accompanying Excel spreadsheet.)

Supplementary Table 7. Phenotypic SNP analysis. The associated gene, chromosome, position, rs tag and observed alleles are shown. For each studied individual, the number of reads supporting each allele is shown.

### Supplementary Text 1: Extended Archaeological Information

#### Levänluhta

The Levänluhta site is located in the Isokyrö municipality at the southern Ostrobothnia region of Western Finland. The site represents a rarely observed case of lake burials, and is one of the most studied archaeological sites in Finland. Levänluhta context has been dated to the Iron Age in Finland (400-800 CE) via some prestige artifacts assumed to have served as grave goods. The skeletal remains are, while numerous and well preserved, also anatomically disarticulated due to the gradual transition of the original lake environment to a marshland, and subsequent ditching and ploughing of the soil for agricultural use over the centuries. The remains of approximately 100 individuals are recognized from the cemetery to date.
The archaeological excavations were carried out by Oscar Rancken in 1886, A.M. Tallgren and Alfred Hackman from 1912 to 1913, Aarni Erä-Esko from the National Board of Antiquities from 1982 to 1984[1], followed by an archaeological survey of both Levänluhta and the immediate area around it in 2014. A comprehensive osteological analysis was reported by Tarja Formisto in 1993. The human remains are under the care of National Board of Antiquities, and stored currently at the National Museum of Finland.

#### Chalmny Varre

The Chalmny-Varre Saami cemetery, associated with the two seasonal settlements of nomadic Kamensk Saami in the 18$^{th}$ century, is located on a small island in the middle flow of Ponoy River (center of Kola Peninsula). The burials have characteristics of Christian graves, combined with old traditional Saami rituals, such as birch cork pieces placed in the graves and masks (lichiny) of deceased carved on wooden crosses. Archaeological dating is confirmed by artifact findings from the graves and information on the wooden crosses marking the graves. The excavation, including an anthropological investigation, was organised by the Institute of Ethnography of N.N. Miklukho-Maclay Academy of Science of the USSR in the year 1976.

#### Bolshoy Oleniy Ostrov

Bolshoy Oleniy Ostrov (Great Reindeer Island), situated in the Kola Bay of the Barents Sea and separated from the mainland by Yekarerininsky Island and two straits, harbors the ancient cemetery of an unknown Early Metal Age culture. The preservation of artifacts made from bone and antler, wooden structures, as well as human remains is remarkable for the location and age this site represents. Altogether 19 skeletons of adults and children have been recognized from both single and collective burials of the site, together with more than 250 artifacts. Archaeological surveys and excavations at the location were performed by G.D. Richter and S.F. Yegorov in 1925, by A.V. Schmidt on the USSR Academy of Science Kola Expedition in 1928, by N.N. Gurina in 1947–1948, as a part of Kola Expedition from the Leningrad Department of the Institute of Archaeology of the Academy of

Sciences, and by V.Y. Shumkin from the same institute, later named as the Institute for the History of Material Culture Russian Academy of Sciences (RAS), starting from 1998-1999 and continuing in 2001-1004. Apart from these excavations, approximately 25 burials were revealed in 1934 during the construction of fortifications. Four finds are known to have been stored by the USSR Academy of Sciences at the time, but the location of all other remains from this instance is unknown. Part of the cemetery was never excavated and has possibly been destroyed by erosion. Morphological analyses, largely concentrated on cranial characteristics, have been performed by S.D. Sinitsyn in 1930, and V.P. Yakmov in 1953 and V.G. Moiseyev and V.I. Khartanovich in 2012[2]. Radiocarbon dates are provided by Moiseyev and Khartanovich in their 2012 study, placing the site in middle to the late 2nd millennium BC. The human remains are stored at Peter the Great Museum of Anthropology and Ethnography (Kunstkamera) RAS in St. Petersburg, with the exception of burial 13, the remains of which are admitted to the Historical Museum in Polarnyi, Murmansk Oblast.[2]

1. Wessman, A. Levänluhta. A place of punishment, sacrifice or just a common cemetery? *Fennoscandia archaeologica* **XXVI,** 81–105 (2009).
2. Moiseyev, V. G. & Khartanovich, V. I. Early Metal Age crania from Bolshoy Oleniy Island, Barents Sea. *Archaeology, Ethnology and Anthropology of Eurasia* **40,** 145–154 (2012).
